# Mutualisms as engines for rapid adaptation: *Rhizobium* evolution facilitates plant drought resistance

**DOI:** 10.64898/2026.07.14.738479

**Authors:** Kevin D. Ricks, Chase Schwarz, Meghan Blaszynski, Destiny Gonzalez, Jennifer A. Lau, Katy D. Heath, Anthony C. Yannarell

**Affiliations:** University of Toronto, Department of Ecology and Evolutionary Biology, Toronto, ON, M5S 3B2, Canada; University of Illinois Urbana-Champaign, Program in Ecology, Evolution, and Conservation Biology, Urbana, IL, 61801, USA; University of Illinois Urbana-Champaign, Department of Plant Biology, Urbana, IL, 61801, USA; Roswell Park Comprehensive Cancer Center, Department of Cancer Genetics and Genomics, Buffalo, NY, 14203, USA; Indiana University, Department of Biology, Bloomington, IN, 47405, USA; University of Illinois Urbana-Champaign, Carl R. Woese Institute for Genomic Biology, Urbana, IL, 61801, USA; University of Illinois Urbana-Champaign, Department of Microbiology, Urbana, IL, 61801, USA; University of Illinois Urbana-Champaign, Department of Natural Resources and Environmental Sciences, Urbana, IL, 61801, USA

**Author notes:** Co-first authors (Kevin D. Ricks, Chase Schwarz).

**Keywords:** experimental evolution, global change, mutualism evolution, host-microbe interactions, legume- rhizobia symbiosis

## Abstract

Mutualisms play critical roles in organismal stress tolerance; yet environmental stressors may simultaneously alter the evolution of the mutualism itself. Stress may have particularly strong impacts on the evolution of microbial mutualists due to their capacity for rapid genetic change. Here we used a legume-rhizobia mutualism, in which plants exchange carbon for symbiotically fixed nitrogen, to evaluate how mutualisms evolve in response to stressors. We experimentally evolved populations of *Rhizobium leguminasarum* in a full factorial design, manipulating drought and nitrogen. We quantified genomic changes in *Rhizobium* populations as well as their quality as partners with their plant host, *Trifolium repens*. Drought selected for context-dependent stress benefits to the host; drought-adapted *Rhizobium* strains provided increased benefits to the host under drought, but fewer benefits to the host in well-watered environments. Conversely, nitrogen fertilization selected for decreased *Rhizobium* partner quality. Comparative genomics indicated that selection on standing structural variants along the symbiotic plasmid may underpin these drought benefits, specifically along genes associated with desiccation tolerance. These results suggest that stress can expand the benefits rhizobia provide to their legume hosts beyond nitrogen fixation, with rapid symbiont evolution an engine in promoting adaptive plant phenotypes.

**SIGNIFICANCE STATEMENT:** While mutualisms are fundamental to ecosystem functioning, they are often assumed to break down under environmental stress. We showed the opposite can happen. Here, we factorially manipulated drought stress and nitrogen fertilization and experimentally evolved a model legume-rhizobia mutualism, where rhizobial-bacteria trade nitrogen for plant carbon. When droughted, the symbiotic bacteria rapidly evolved new traits that made them better partners under drought, improving plant growth within just a few generations. Evolution occurred mainly on a specialized bacterial plasmid, a mobile piece of DNA distinct from the main genome. By contrast, added nitrogen fertilizer led to decreases in bacterial benefits. These findings suggest that mutualisms are not simply fragile in the face of global change, but can expand mutualistic benefits to buffer partners against stress.

**CLASSIFICATION:** Biological Sciences, Evolution

## INTRODUCTION

While mutualisms provide a myriad of benefits that support organismal health (1), these benefits are often context-dependent and can therefore be sensitive to environmental fluctuations (2–4). Indeed, shifts in mutualist benefits due to abiotic stress are well-documented (5–7). However, the resulting evolutionary responses remain largely unresolved, despite their potential for cascading fitness effects and long-term eco-evolutionary feedbacks between partners (1, 8). With rapidly shifting environments, understanding how interactive, and potentially conflicting, environmental stressors alter the trajectory of mutualism evolution will be essential in predicting organismal stress responses in a changing world.

Environmental stressors could drive mutualism evolution towards two potential outcomes: destabilizing the interaction and increasing stress or, conversely, driving evolutionary innovations that buffer both partners against stress. On the one hand, the sensitivity of these interactions might exacerbate the effects of global change by favoring maladaptive or uncooperative mutualist partners, particularly if the mechanisms that normally stabilize mutualism break down (5, 9, 10). Furthermore, a partner’s dependency on the interaction may generate a mutualistic “two-body problem”, wherein limitations on one partner inherently limit the other (11, 12). These disruptions may drive the decline or even collapse of mutualism. On the other hand, phylogenetic evidence suggests that mutualisms can be highly stable interactions (13, 14) and engines of diversification over evolutionary time (15, 16). Thus the evolution of mutualistic partners might instead facilitate rapid innovations that promote adaptation to novel environments, making organisms robust to environmental perturbation (17). Given the central role of mutualisms across ecosystem functioning, including carbon and nutrient cycling, seed dispersal, and plant pollination (1), these alternative outcomes have significant implications for ecosystem resilience.

Eukaryotic organisms host diverse microbial communities, including many mutualistic strains, that are increasingly recognized to play fundamental roles in host development, phenotype, and survival (18, 19). These host-microbe mutualisms provide powerful experimental systems for examining the impact of global change on mutualism evolution and stability. Due to their short generation times and large, genetically-diverse populations (20), these microbial symbionts can undergo significant evolutionary shifts within only a few host generations (21).

Moreover, many microbes possess modular genomes with mobile genetic elements (MGEs) such as plasmids and viruses, which can evolve rapidly and follow evolutionary trajectories independent of the bacterial chromosome (17). For example, mutualistic bacteria frequently carry the genes essential for symbiosis on plasmids, which are hotspots for genetic diversity and facilitate the rapid movement of these key genes within populations (18). Experimentally evolving microbial mutualists is a tractable approach for examining the genetic mechanisms by which the vast diversity in microbial genomes and their MGEs contribute to organismal stress adaptation (22).

Here we use the mutualism between leguminous plants and nitrogen-fixing rhizobial bacteria to understand how mutualist stress-adaptation shapes host resilience. Rhizobia are a model for mutualism evolution, easily culturable, have well understood genomic architecture with key symbiotic traits hosted on MGEs, and can rapidly evolve in response to environmental conditions. Rhizobial bacteria specialize in infecting the root nodules of leguminous plants, wherein they fix atmospheric nitrogen (N) into biologically-available forms in exchange for plant sugars (23). This symbiosis is critical in global ecosystem functioning, as a primary driver of N cycling (24).

We experimentally manipulated drought and nitrogen to evaluate their impact on rhizobia evolution. We maintained mixed populations of *Rhizobium leguminosarum* strains in a full factorial design, manipulating drought, N-fertilization, and the presence of the host plant *Trifolium repens*, which allowed us to evaluate whether evolutionary response of microbial mutualists to these environments is mediated by their plant partners. Drought is a ubiquitous abiotic stressor intensified by global change, and decreases fitness of both plant and rhizobia partners (28). Nitrogen (N) fertilization, while not a physiological stress *per se*, can fundamentally disrupt resource exchange and thus destabilize the mutualism, as plant partners decrease investment in mutualistic interactions in response to abiotic sources of N (8, 29).

Evolving microbial mutualists in response to two, potentially orthogonal, stressors allows us to examine the additive effects of these historic perturbations on mutualism function and thus the benefits to hosts. By re-isolating single strains after experimental evolution and resolving reference-quality whole genome sequences, we identify the genomic underpinnings of this adaptation.

## RESULTS

### *Rhizobium* stress adaptation generates context-dependent *Rhizobium* partner quality

We isolated 320 descendant strains from replicate populations that had evolved in 8 selection environments (drought x N x host presence). We then used single strain inoculations of these evolved strains in a “test phase” to characterize shifts in the quality of *Rhizobium* partners (partner quality, i.e. *Rhizobium* impacts on plant growth) in both wet and dry conditions. Because our starting *Rhizobium* population was a mix of 28 strains with varying partner quality across wet and dry environments (Fig. S1), differential phenotypes between treatments can be a result of both shifts in strain frequencies as well as introduced *de novo* mutations.

Rhizobia rapidly evolved in response to dry and wet soil moisture conditions in ways that improved plant fitness under those growth conditions; however, these increases in *Rhizobium* quality were only observed when strains had evolved in the presence of a host plant and unfertilized soils (Fig. 1, S2–S4, Tables S1–S2). In other words, *Rhizobium* partner quality was maximized when strains were evolved 1) with a plant partner, 2) without N-fertilizer, and 3) in a watering history that matched the contemporary watering condition of the test phase (e.g., wet history under wet conditions, drought history under droughted conditions). Partner quality in these evolved strains was highly contingent on the presence of the host during the selection phase, with minimal variation between treatments when rhizobia had evolved in the absence of a plant (Fig. 1). These shifts in partner quality between water histories are not only the result of evolutionary changes in N fixation; similar patterns were observed when plants received supplemental N fertilization in a small follow-up experiment (Fig. S5). However there may be some variation in fixation between nitrogen histories as plants inoculated with N-fertilizer adapted strains had marginally lower plant tissue nitrogen content (Fig. S4).

**Fig. 1:**
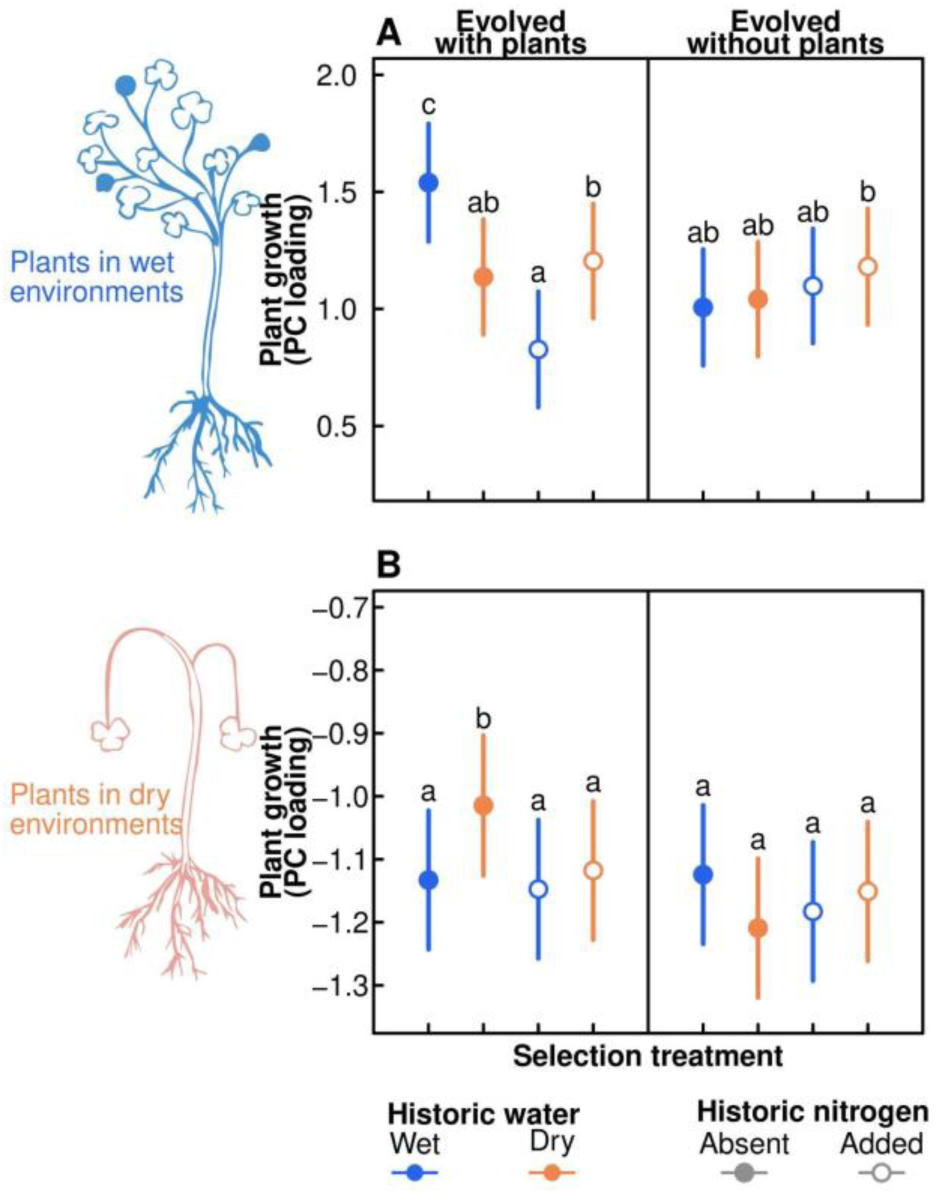
*Rhizobium* populations that had evolved in wet conditions benefit plant growth in wet environments **(A)** and rhizobium population that evolved in dry conditions benefit plant growth in dry environments **(B)**, but only when rhizobia evolved in the presence of plant hosts and the absence of N-fertilization. Shown are estimated marginal means (with 95% confidence intervals) of partner quality (PCA including biomass, height, and leaf number) from a common garden in which plants were inoculated with one of 320 descendant strains isolated from *Rhizobium* populations evolved for four plant generations in factorial combinations of drought, N fertilization, and host presence. Letters represent significant differences according to *post hoc* Tukey tests. Note the separate y-axis scales in top and bottom panels.

### Drought treatments select on *Rhizobium* standing variation

To identify the bacterial genetic mechanisms underlying the evolved partner quality variation, we sequenced 160 of the strains that evolved with a plant partner, then mapped these genomes back to previous assemblies for the 28 ancestral lineages (17). This approach allowed us to assess the contribution of shifts in standing genetic variation, any large-scale plasmid HGT, and the emergence of *de novo* mutations to the observed phenotypes.

Our drought and nitrogen treatments selected for significantly different sets of *Rhizobium* strains, with different re-isolation frequencies of bacterial ancestral lineages between treatments (i.e., different ancestral lineages became dominant in dry versus wet treatments; Fig. 2A & S6A, Table S3) indicating environment-dependent selection on standing variation. Model selection strongly suggested that ancestor isolation frequencies were best explained by the ancestral strain’s partner quality as measured in the matched watering environment, as opposed to partner quality measured in the mismatched watering environment (Table S4). For example, ancestral lineages with the highest benefits to plant growth in wet conditions (measured in an independent experiment) became dominant in wet evolutionary treatments (Fig. 2B), while those that provided the highest benefits in drought conditions became dominant in dry treatments (Fig. 2C).

**Fig. 2:**
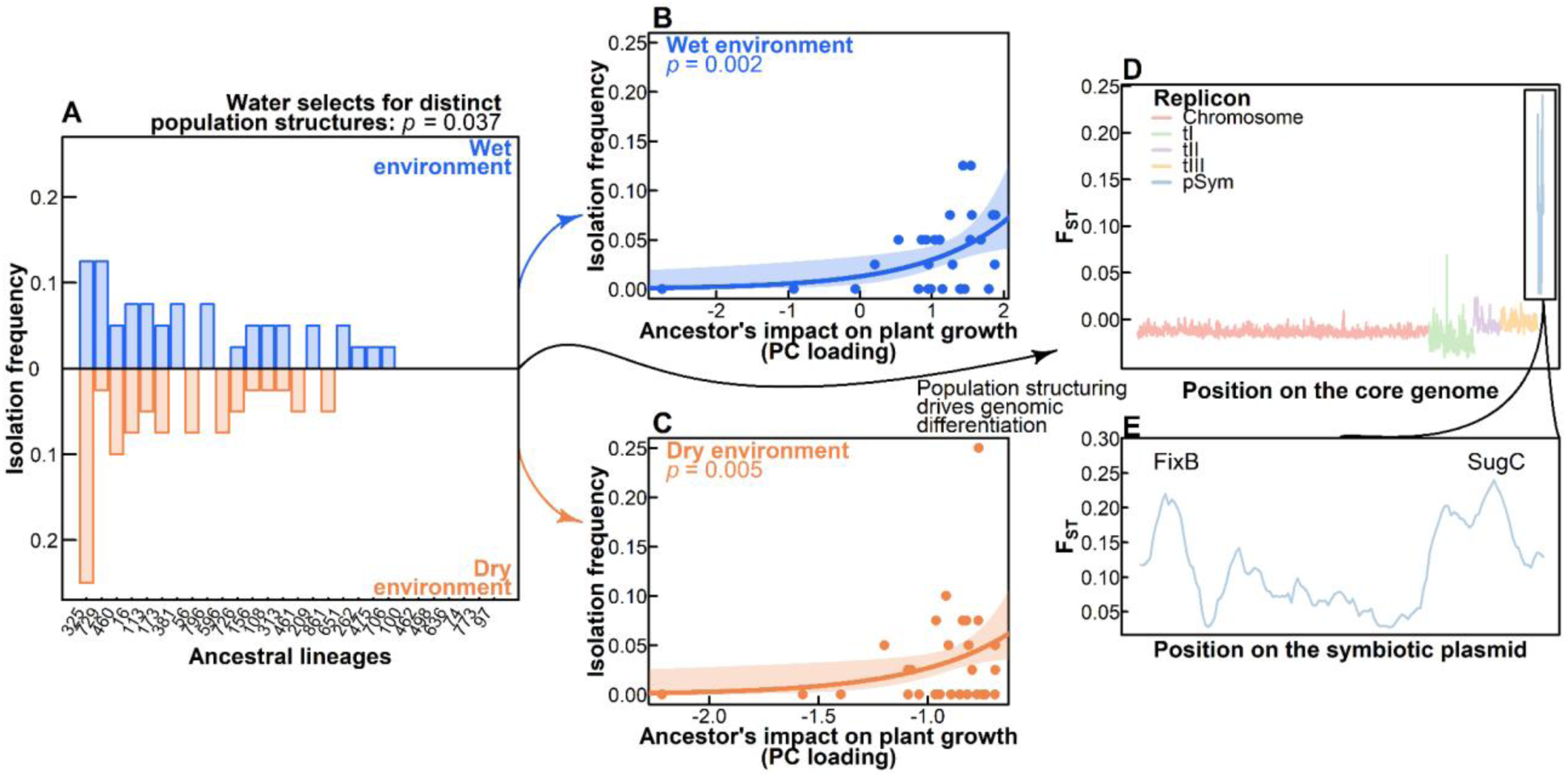
Watering history shapes *Rhizobium* evolution through selection on standing variation in the population. We display **(A**) the isolation frequencies of the original 28 ancestral lineages in the two watering treatments evolved without nitrogen fertilization, including a *p* value representing a χ^2^ characterizing differentiation between populations. We relate ancestral frequency to benefits to impact on plant growth **(B, C)**, correlating ancestral isolation frequencies in wet environments with ancestral partner quality in wet, and ancestral isolation frequencies in dry environments with ancestral partner quality in dry. We include *p* values in the top left for these models describing this correlation. Variation in strain frequencies underlies genomic differentiation (F_ST_) between populations **(D, E)**. We highlight two genes along the symbiotic plasmid (pSym) that pass a 0.2 F_ST_ threshold, representing regions of significant differentiation.

Shifts in strain frequencies between watering treatments was accompanied by significant genomic differentiation, primarily localized to the symbiosis plasmid (pSym). An F_ST_ sliding window analysis based on the core genome identified significant genomic divergence between wet and dry treatments occurring exclusively along the pSym (Fig. 2D–E). The regions of highest differentiation overlapped with *fixB* (a component of the nitrogen fixation pathway) and *sugC* (an osmolyte transport gene), with F_ST_ values significantly differing from simulated evolution under drift (Fig. S7). Similarly, analysis of gene presence/absence variation using Fisher’s exact test identified 40 genes with significantly different frequencies between the wet and dry treatments (*p* < 0.05 corrected for false discovery, Table S5), the vast majority of which (68%; 27 of 40) were also located on this replicon.

This pSym-driven genomic divergence was underpinned by selection on distinct plasmid genotypes. Our prior work has indicated that *Rhizobium* carry one of four major pSym types (30), whose evolutionary histories are discordant with the main chromosome (termed A–D, Fig. 3A). The isolation frequencies of these plasmid types significantly differed among selective treatments (Fig. S6B). The dry treatment, for example, was comparatively enriched in strains containing pSym type A and depleted of strains with pSym type C (Fig. 3B). This shift in pSym type was directly correlated with the observed gene presence/absence variation between treatments. For example, the 1-aminocyclopropane-1-carboxylate deaminase gene (*acdS*), notable for its function in plant drought resistance (31, 32) was highly enriched in dry-isolated strains (97% versus 57% in wet-isolated strains; corrected *p* < 0.001). This variation was strongly associated with pSym type, as this gene was ubiquitous across the population with the exception of pSym type C (Fig. S8), the same clade depleted in the drought treatments. We note that this pSym-driven variation was the result of selection on standing variation rather than the horizontal transfer of plasmids during the experiment, as we detected only a single instance of plasmid loss across all sequenced strains. No instances of plasmid exchange were found.

**Fig. 3:**
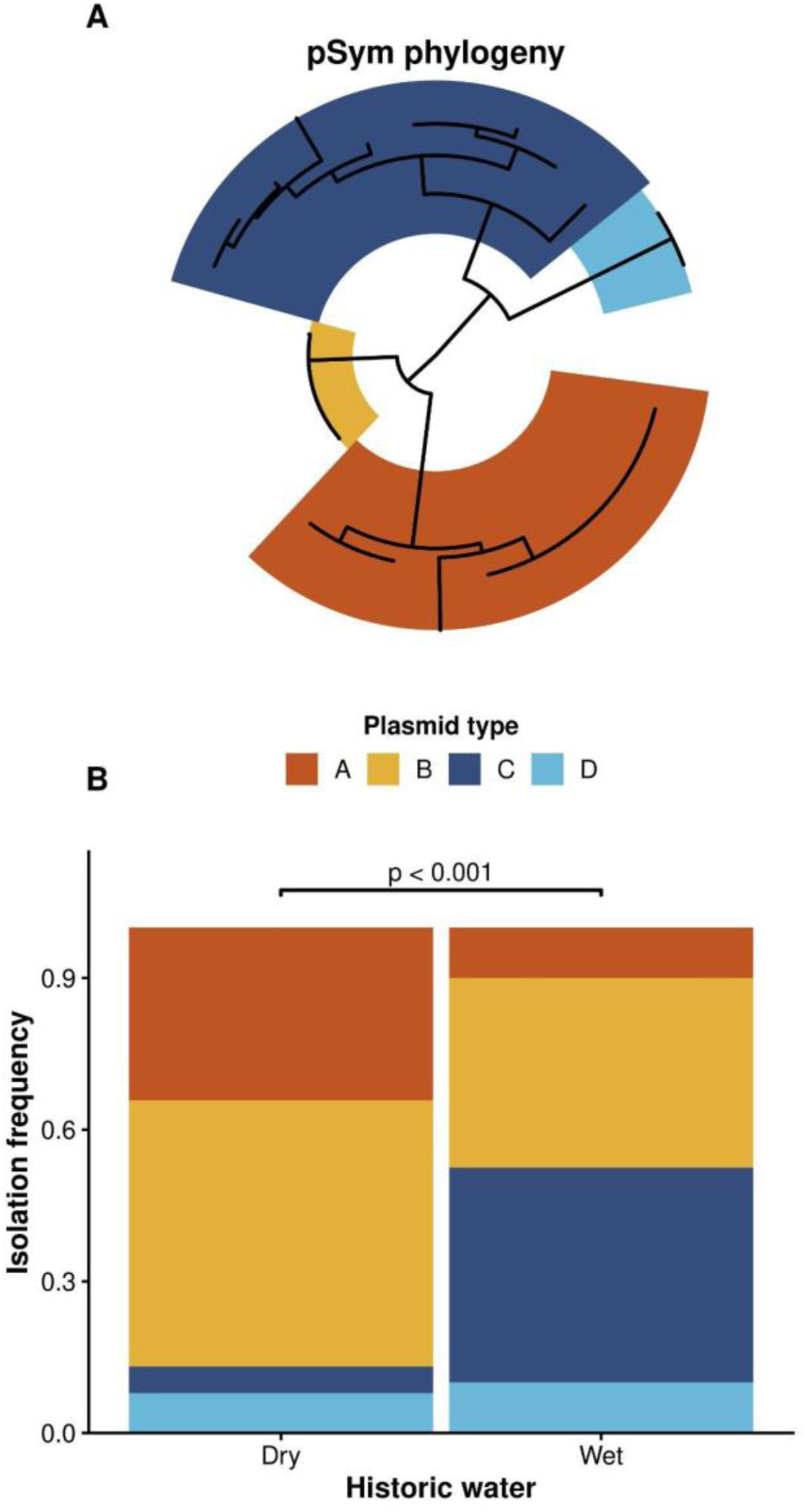
Phylogeny of the pSym structures population differentiation between the wet and dry treatments. Our prior work in this system (30) has identified 4 major types of pSym, structured along phylogeny **(A)**. The isolation frequency of strains between historic watering treatments, under low nitrogen fertilization, is significantly structured by these plasmid type frequencies based on χ^2^ tests **(B)**.

In contrast to the response in the watering treatments, adaptation to N-fertilization treatments yielded little evidence for selection on standing variation. Comparing the subset of strains from wet environments, since N-fertilization had the largest impact on phenotype in wet-adapted populations, we found minimal variation in strain isolation history (Fig. S6) or F_ST_ (Fig. S9) between nitrogen treatments and only a few genes with significant presence/absence variation (7 genes, Table S6).

### Tradeoffs in *Rhizobium* partner quality through *de novo* mutations

Comparing the sequenced evolved strains to their putative ancestors identified a total of 655 single nucleotide polymorphisms (SNPs). Of those, 395 came from two hyper-mutator strains, which we excluded from the remaining analyses. SNPs were not evenly distributed across treatments or replicons (Table S7), generally occurring at the highest rate on the pSym, particularly in the wet strains (e.g., chromosome: 3.35 x 10^-6^, pSym: 1.49 x 10^-6^, Table S8).

While the overall number of SNPs is low, introduced *de novo* variation nonetheless appeared to contribute to the observed variation in plant phenotype. Even after controlling for the ancestral lineage, partner quality was maximized when the contemporary watering environment matched the selective history (Fig. 4A–B, Table S10–S11). Furthermore, we identified subtle but significant tradeoffs in partner quality between wet and dry environments; when controlling for ancestral background, strains that conferred the highest plant growth benefits in droughted environments conferred fewer benefits in wet environments, and vice versa (Fig. 4C; *p* = 0.029).

**Fig. 4:**
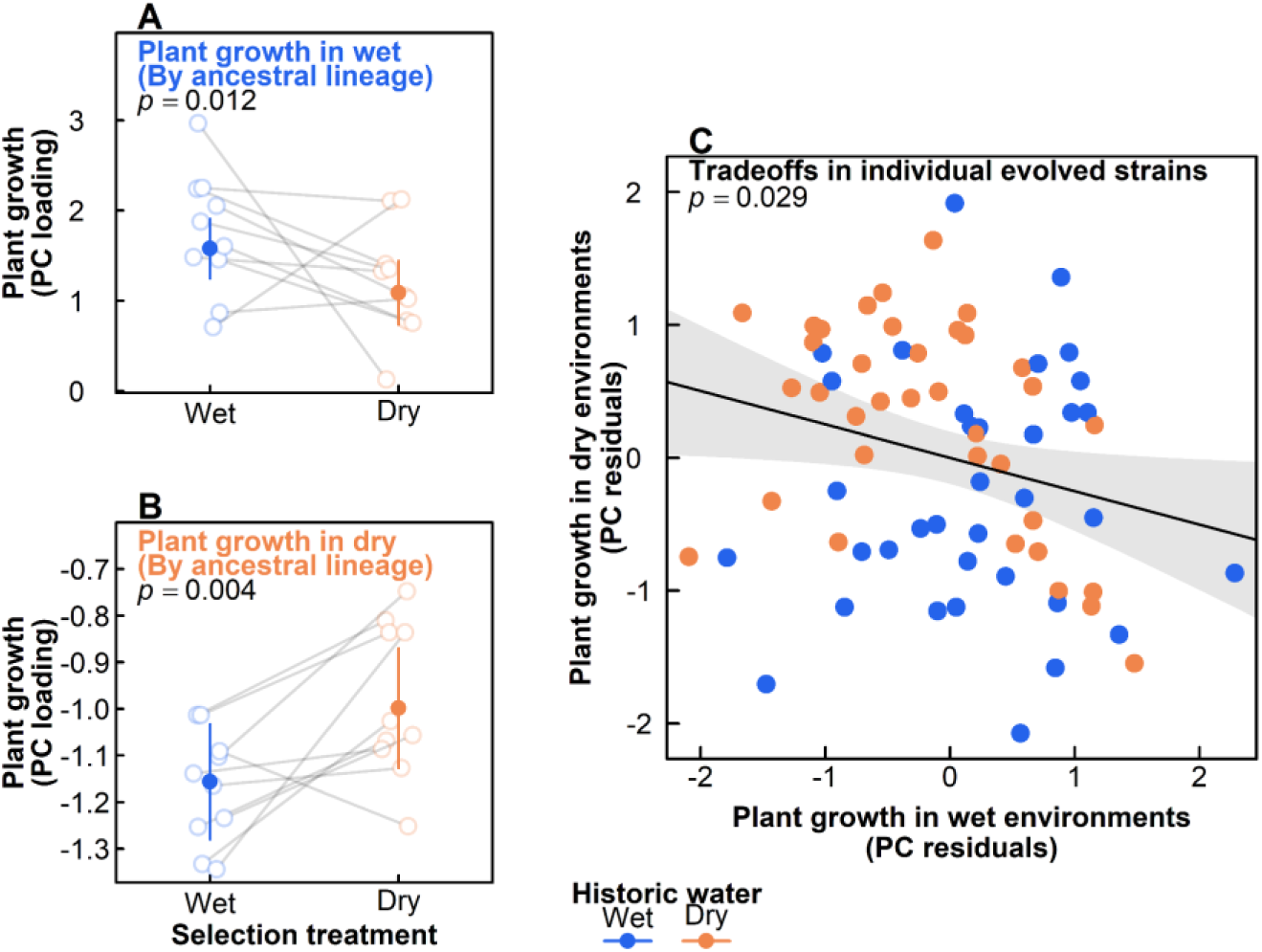
Evolutionary history of the experimentally evolved *Rhizobium* populations impacts their partner quality, even when controlling for ancestral lineage, in both contemporary wet **(A)** or contemporary dry **(B)** watering environments, driving genetic tradeoffs between wet and dry environments. We include only strains evolved in unfertilized nitrogen treatments. Semi-transparent points represent the mean of evolved strains belonging to each ancestral lineage when plants were grown in wet **(A)** or dry **(B)** environments. Solid point represents the group marginal mean while controlling across ancestral lineages, with bars representing 95% confidence intervals generated from the standard error. Points colored by strains’ historical watering treatment: wet as blue, and dry as orange. From the residual variation following accounting for ancestral lineage we find a weak negative correlation between plant growth in wet and dry environments, suggesting that *de novo* mutations further contribute to trade-offs between wet and dry environments **(C)**.

Interestingly, this context-dependent shift in partner quality was correlated with the accumulation of SNPs. Within ancestral lineages, an increase in the number of SNPs among descendant strains was significantly associated with increased partner quality specifically in environments that matched the strain’s selective watering history (Fig. S11). While large structural variations on the pSym drove major population-level shifts, this accumulation of genetic changes between ancestors and their descendants contributed to increased partner quality in matching watering environments.

Similar results were found in comparing N-fertilization treatments within the ancestral lineages. Partner quality within ancestral lineages significantly decreased when evolved with added N-fertilization (Fig. S12, Table S12), and decreased partner quality was associated with an increasing number of SNPs in nitrogen-adapted strains (Fig. S13).

Evaluating SNP distribution based on their association with specific gene annotations, we found both the historic watering and N treatment to be significant predictors (Fig. S10, Table S9). Dry-adapted strains were enriched for SNPs in genes associated with cell membrane, while wet-adapted strains were enriched for SNPs in genes associated with hydrolase activity and DNA binding. High nitrogen-adapted strains were enriched for SNPs in genes associated with metal ion activity and ATP binding. Low nitrogen-adapted strains were conversely enriched for SNPs associated with peptidases, proteolysis, and cytosol (see Supplemental Data for full list of all SNPs and their associated annotations).

## DISCUSSION

A critical, unresolved question is whether mutualisms will collapse under environmental change, or buffer partners against these changes. We found evidence for both outcomes. Nitrogen- fertilization selected for less cooperative *Rhizobium* mutualists, consistent with prior work (8).

Drought, however, selected for rhizobia that maintained plant fitness in their local moisture conditions. Namely, plant growth was maximized when hosts were inoculated with *Rhizobium* evolved in moisture conditions that matched the plant’s current environment. This rapid adaptation of *Rhizobium*, in the space of only a few plant generations, stemmed from selection on the symbiosis plasmid: this plasmid was the hotspot for *de novo* mutations, as well as containing key genes associated with drought resilience which were enriched in the drought treatments. As these drought benefits only emerged when rhizobia evolved with a plant, host presence facilitated this selection. Below we discuss mechanisms of bacterial adaptation and their implications to mutualism evolution.

### Mechanisms of rhizobial drought benefit

Strikingly, *Rhizobium* populations evolved in response to watering treatments in ways that directly benefit hosts in those treatments — i.e., in a direction consistent with local adaptation for the host plants. One mechanism for this rhizobial drought benefit is the evolution of drought-tolerant nitrogen fixation processes, as the fixation enzyme, nitrogenase, is sensitive to stress (33, 34). Such efficiencies could generate the observed drought benefits, as nitrogen availability can alleviate plant drought stress through the maintenance of plant hormones/antioxidants and water use efficiency (35, 36). Our data, however, provided mixed support for this hypothesis. On one hand, differentiation between wet and droughted populations at fixation-associated genes (e.g., *fixB*) highlights potential differential selection on nitrogenase between watering treatments.

However, increased nitrogen fixation efficiency under drought should be associated with increased plant tissue nitrogen content — this was not observed. Furthermore, in a follow-up experiment, the drought-adapted rhizobia provided increased growth benefits even in high-nitrogen environments where benefits from nitrogen fixation should be minimized.

These rhizobial drought benefits may, therefore, have emerged through mechanisms unrelated to nitrogen fixation. For example, we observed significant genomic differentiation between droughted and wet populations along the symbiosis plasmid, notably at sugC, an osmolyte transport gene associated with trehalose production. Bacterial trehalose production can increase plant osmotic tolerance (37), acting as a stress-response signaling molecule (38) as well as an osmoprotectant, maintaining cell membrane integrity (39). Similarly, the drought-adapted population was enriched in strains with *acdS* genes (Fig. S8). While ethylene is an important plant hormone, high concentrations produced during stress can inhibit growth and accelerate senescence (40). Bacterial *acdS* can alleviate plant stress through degradation of the ethylene precursor (32). In addition, we identified a number of introduced mutations that may be contributing here. Relative to their ancestors, dry-adapted descendant strains were enriched in SNPs associated with cell membrane function, which could be associated with bacterial osmotic regulation and desiccation tolerance. These results suggest diverse pathways by which rhizobia facilitate plant drought stress and offer targets for future exploration.

### Local adaptation and expansion of mutualistic benefits

With the emergence of these novel benefits, drought may be selecting for an expansion of the legume-rhizobia mutualism beyond nitrogen fixation alone. While prior case studies have identified specific rhizobial traits that benefit droughted plants, such as small molecule production (32), our results provide direct experimental evidence that rhizobia can evolve such benefits over rapid timeframes in direct response to drought. These results run counter to predictions that mutualisms simply collapse under stress (5). Instead, rapid rhizobial evolution appears to generate stress-adapted phenotypes in the host plant. These phenotypes could emerge as a result of rhizobial dependencies on the plant for carbon resources, generating selective pressure for rhizobial traits that facilitate plant drought tolerance in order to maintain access to these resources.

These context-dependent benefits may explain the variability in mutualistic partner quality. While theory predicts selection should minimize partner quality variation and select for solely high-quality partners (41), natural populations host enormous variation (42). The emergent tradeoffs partner quality in our evolved strains suggest some of the partner quality variation may be the result of heterogeneous environments selecting for context-dependent, locally adaptive partners. Localization of these benefits on the pSym may facilitate both the maintenance of these traits and their rapid spread, a point we return to below. Consistent with this hypothesis, while the ancestral strains used herein were chosen as seemingly random subset of strains from a natural clover population, these drought-tolerance traits were pre-existing here. Isolated originally from an early successional grassland with a history of periodic droughts (43), these environments may have facilitated past genomic innovations generating the drought-adaptive plasmids, and maintained in the population through seasonal water variability..

### Nitrogen fertilization erodes mutualism quality

In contrast to the watering environment, N-fertilization selected for reduced rhizobial quality. This reduction may be the result of N-fertilization disrupting the primary axis of resource exchange in this mutualism. While nitrogen fixation is metabolically expensive, mechanisms such as plant partner choice and fitness feedbacks maintain and stabilize these mutualistic traits (44, 45); added nitrogen, however, may undercut these, as the benefit rhizobia provide to the plant host is minimized and plants invest less in their rhizobium partners (29). Interestingly, in contrast to past work (30), N-fertilization did not alter the isolation frequency of ancestral lineages as the drought treatment did, instead selecting for mutations at sites associated with specific functions, such as metal ion activity and ATP binding. As the nitrogenase enzyme has high energy demands while requiring specific enzymatic cofactors (metal ions), loss-of-function mutations in these pathways may provide a consistent advantage by shutting down costly machinery. Consistent with this, tissue N marginally decreased when plants were provided with rhizobia evolved with added N.

### Genomic underpinnings of rapid symbiont adaptation

Why could these microbial symbionts evolve so quickly? Our work points to the pSym as a selective hotspot underlying the rapid emergence of drought phenotypes. Indeed, this plasmid was both a region of high standing genetic diversity and the primary site of introduced mutation for drought-related traits. Plasmids may be natural locations for drought-tolerance traits given their propensity for rapid evolution; they are frequently enriched in genes mediating adaptation to extreme environments, as such genes can be quickly gained or lost from a population under stress (46, 47). Plasmids also recombine at higher rates and accumulate *de novo* mutations faster than the chromosome (48), which may generate and sustain the observed increased genetic diversity. While the pSym carries the core nitrogen-fixation genes, thought to be co-located on a single plasmid because they function together during the fixation process (26), accessory genes can also be found here (30). The drought-tolerance loci identified here, including *acdS* and trehalose biosynthesis, may represent another class of accessory trait accumulating on this already gene-rich, rapidly evolving replicon. As regions of diversification, plasmids may be playgrounds for genomic innovations, producing reservoirs of genetic diversity that can quickly move through the population and underpin adaptation.

While we’ve identified some genomic drivers of the evolved phenotypes, there remained substantial variation within lineages with no detected SNPs. Epigenetic modifications could be playing a critical role in driving this variation. In bacteria, epigenetic modifications, notably DNA methylation, can rapidly alter transcription. While many epigenetic marks are transient, specific methylation states can regulate phase variation and be vertically inherited across multiple generations (49, 50). Such modifications can be stable across replication cycles (49), representing a mechanism for rapidly reshaping bacterial phenotype here. Future work may consider approaches for investigating transcriptional variation in these systems.

## CONCLUSIONS

Our results suggest that environmental changes can drive the evolution of symbiont populations toward different outcomes depending on the type of environmental change. While N-fertilization eroded *Rhizobium* partner quality, drought selected for symbionts that promote plant drought tolerance. Genetic variation along the symbiosis plasmid, including new mutations and selection on pre-existing variation drove the observed rapid evolution. Moreover, this plasmid hosted genes for both nitrogen-fixation and drought-stress tolerance. If environmental variation generates and maintains this and other context-dependent functions, partner quality may be shaped not only by nitrogen exchange, but by additional traits tied to environmental stress tolerance. Overall, our work suggests mutualisms can be robust against stress, and may in fact, be engines for introducing adaptive phenotypes in their hosts.

## MATERIAL & METHODS

### Experimental Evolution

We evolved populations of *Rhizobium leguminosarum* comprising 28 ancestor strains (isolated from Kellogg Biological Station (8)) over four plant generations. We used a full factorial design crossing water (wet vs. dry), fertilization (no nitrogen vs. nitrogen added), and host presence (plant vs. bare soil) across 120 soil mesocosms (n=15 replicates per treatment combination). In the first generation, pots were inoculated with an equal mixture of all 28 ancestor strains.

Moisture treatments were maintained through watering pots every 2 days to either saturation (wet) or 14% gravimetric water content (dry). After each 10-week generation, plants were cut back, and lay fallow for three weeks to release rhizobia into the soil from nodules before the next generation was sown.

### Rhizobia Isolation and Phenotyping

Following the selection phase, we isolated rhizobia from the soil by using a “trap plant”, growing plants under wet, no nitrogen conditions, following prior work (8). Ideally we would isolate rhizobia directly from the soil to prevent biases introduced from selection in this phase. This, however, can be prohibitively inefficient in recovering rhizobia relative to other soil bacteria.

With our approach, we isolated directly from the nodule, recovering 446 descendant strains, from which 320 (40 strains per drought x fertilization x plant treatment combination) were selected for phenotyping.

To assess strain partner quality, we inoculated each strain onto three replicate *T. repens* seedlings in both wet and dry environments (1,960 plants, as well as sterile controls). After eight weeks in the greenhouse, we measured above- and belowground biomass, plant height, leaf number, and root nodule counts. We analyzed a subset of plants (n=160) from the dry environments for tissue N and δ^15^N to characterize nitrogen fixation capacity.

### Genomic Sequencing

We performed PacBio HiFi long-read sequencing on a subset of evolved strains. While long-read sequencing has higher costs, we chose this as other work has suggested that genomic rearrangement can underpin *Rhizobium* evolution (51). Due to these costs, of the 320 strains collected we sequenced only those evolved with a plant (160 strains). Genomes were assembled *de novo* and mapped to the 28 ancestral references based on highest average nucleotide identity. We identified single nucleotide polymorphisms (SNPs) by comparing evolved strains to their predicted ancestral references with standard SNP-calling pipelines.

### Statistical Analysis

#### Greenhouse phenotypes

We analyzed plant growth traits using linear mixed-effects models in each contemporary watering environment. Fixed effects included historic moisture, historic nitrogen, and historic plant presence; greenhouse block and strain were random effects. We additionally compressed the aboveground traits, representing plant growth and therefore good proxies for *Rhizobium* partner quality, to a principal component axis (PCA) to broadly evaluate variation along a single axis. This was paired with a Redundancy Analysis (RDA) as a multivariate approach to examine how this suite of traits were impacted by these variables.

#### Population Genetics

Of the 160 sequenced strains, 155 strains could all be mapped back to 22 of our 28 ancestral strains. One strain was a contaminant and clearly not related to any of our ancestral strains, while the other 4 had low sequencing depth.

We compared ancestral re-isolation frequencies across treatments using χ^2^ tests. As our prior work has identified 4 major clades in pSym phylogeny (30), in parallel we used χ^2^ to compare pSym clade frequency. We additionally correlated strain quality with isolation frequency using generalized linear mixed effects models with a poisson distribution. Genomic differentiation (F_ST_) was calculated between moisture and nitrogen treatments, as well as gene presence/absence variation using Fisher’s Exact test.

We evaluated the contribution of *de novo* mutations to observed phenotypes by categorizing gene functions in which SNPs occurred into Gene Ontology (GO) terms. Using multivariate and univariate models, we predicted SNP distribution across the suite of GO terms based on their historic selective treatment.

## ACKNOWLEDGEMENTS

This research is a contribution of the GEMS Biology Integration Institute, funded by the National Science Foundation DBI Integration Institutes Program, Award #2022049. It was additionally supported by the Cooperative State Research, Education, and Extension Service, US Department of Agriculture, under project number ILLU 875-952, as well as by the School of Integrative Biology and the Graduate College at the University of Illinois Urbana-Champaign. We thank Frederickson lab members at University of Toronto for thoughtful feedback on early drafts of this manuscript. *Rhizobium* strains used in this experiment were isolated from the KBS LTER, which is supported by the NSF Long-term Ecological Research Program (DEB 2224712) at the Kellogg Biological Station and by Michigan State University AgBioResearch.

## DATA AND CODE AVAILABILITY

All sequence data is available at the NCBI Genome and SRA databases, https://www.ncbi.nlm.nih.gov/bioproject/PRJNA1294393.

## AUTHOR CONTRIBUTION

KDR, KDH, JAL, & ACY conceived the initial framework for the study. KDR conducted the initial greenhouse experiments, while MB and DG conducted the nitrogen-addition experiment.

CS was responsible for all bioinformatic work. KDR and CS wrote the first draft of the manuscript, and all authors contributed to subsequent revisions.

## SUPPLEMENTAL METHODS

### Rhizobium Experimental Evolution

#### Selection Phase (Generations 1–4)

We started this experiment using a collection of 28 strains of *Rhizobium leguminosarum* that were previously isolated from the Kellogg Biological Station (1). With these strains, we established an evolution experiment to investigate how environmental perturbations influence the stability of the legume-rhizobia mutualism. The selection regimes constituted a fully factorial crossed design of three main factors: moisture (wet or dry conditions), nitrogen (nitrogen addition or no nitrogen addition), and presence of a plant partner (grown with *Trifolium repens* plants present or only in bare soil pots).

For the first generation, we established soil mesocosms by filling pots with 430 g of a locally produced “root-wash” soil mix: equal parts of calcined clay, torpedo sand, and field soil (UIUC Plant Care Facility, https://pcf.aces.illinois.edu/soil-mixes/). We prepared 15 replicate pots in each of the eight selection treatments, for a total of 120 pots. We surface-sterilized white clover seeds (*Trifolium repens* L., seed sourced from Ernst Seeds, Meadville, PA, USA) by vortexing them in a 3% sodium hypochlorite and 0.01% Tween-20 solution for 2 minutes, followed by a thorough rinsing in sterile water. We planted approximately 100 seeds into each of the 60 “plant partner present” pots, leaving the other 60 pots with bare soil only. All pots were watered to saturation daily for 1 week after sowing to facilitate successful germination and plant establishment, after which the moisture treatments were imposed: in the wet treatment, pots were watered to saturation every 2 days while in the dry treatments, we weighed pots and watered to 14% gravimetric water content every 2 days. We chose this water content target for the dry treatment as a pilot demonstrated it represented a significant stress compared to the well-watered control, while still allowing for plant survival. We additionally amended mesocosms designated to the ‘nitrogen addition’ treatment with 0.03 g of ammonium nitrate at the start of each generation. This fertilization rate was chosen to be similar to the nitrogen fertilization levels in agronomic settings (2) and has been associated with lower-quality rhizobia partners (1).

We prepared the initial ancestor inoculum for the first generation, by growing each ancestor strain separately in liquid TY medium (0.5% tryptone, 0.3% yeast extract, and 10 mM CaC1_2_) at 30°C for 2 days. We then diluted these cultures to a standard optical density (OD_600_) of 0.1, pooling all 28 ancestor strains into a common inoculum of which 10 mL was added to each mesocosm pot 7 days after sowing.

Mesocosms were grown in the greenhouse for 10 weeks on a 26°C/24°C day/night schedule, supplemented with 14 h of daily light. After 10 weeks, we concluded the generation by pruning back plant shoots and allowed all pots to sit fallow for three additional weeks. This step allowed the senescing root nodules to release rhizobia cells back into the soil (3). We then re-established each of our eight treatment conditions for each mesocosm by sowing surface-sterilized seeds, adjusting the watering regime and supplementing with nitrogen as appropriate per their assigned treatment. We did not re-inoculate with ancestor strains, instead relying on the rhizobia that persisted in the soil or were released from senescing root nodules to establish the next generation. This process was repeated for a total of four plant generations under the selection treatments.

#### Isolation

After four plant generations under selection, we isolated rhizobia from all mesocosms. Because the selection phase included treatments without any plants, we included a fifth “trap plant” generation in the greenhouse to allow us to isolate rhizobia from root nodules. While this introduces potentially biases, specifically selecting for strains that can associated with a plant, this was a necessary choice as isolation directly from the soil can be very difficult.

We prepared new pots filled with sterile soil and 100 surface sterilized clover seeds as above. We then prepared soil slurries from our selection phase pots using 30 g of soil from each mesocosm diluted in 200 mL of a sterile 0.9% NaCl solution. We inoculated 5 mL of these soil slurries into the trap plant pots one week after sowing. We watered the trap plants to saturation every 2 days, under identical climate conditions as the prior experiment, for 5 weeks in the greenhouse, after which they were harvested for nodule isolation.

From each pot, we randomly selected six trap plants for isolation. We chose for isolation those nodules closest to various predetermined & haphazard locations on the root system to avoid biasing the sampling by nodule size. The selected nodules were surface sterilized with bleach and ethanol, squashed to expose rhizobia, and plated onto TY agar. Strains were serially plated to isolate individual colonies and then grown to a high density in liquid TY to be frozen in a 75% TY and 25% glycerol solution at -80°C for future use. We verified the identity of these rhizobia using sequencing. Specifically, we PCR amplified the 16S region using 8F and 1492R primers and submitted samples for both forward and reverse Sanger sequencing at the Roy J. Carver Biotechnology Center (Urbana, IL, USA). In total, we recovered 446 “evolved” *Rhizobium* strains that had passed through the selection phase of the experiment.

#### Assessing evolved rhizobia impact on plant growth

We evaluated the partner quality of the evolved strains. From each of the eight treatments, we randomly selected 40 strains spread as evenly as possible from among the replicate populations. We filled sterilized “cone-tainer” pots with autoclaved sterilized root-wash soil mix. For each strain, we grew 3 replicate plants under well-watered conditions and 3 replicate plants under droughted conditions. We also grew 40 uninoculated plants under watered or droughted conditions as negative controls, for a total of 1960 plants (2 watering treatments x 320 strains x 3 replicates + 40 control (20 wet and 20 dry plants)). We sowed approximately 10 surface sterilized clover seeds in each pot, watered pots to saturation every other day, and then thinned each pot to a single seedling. Seven days after the sowing, we inoculated plants with their designated rhizobia strain and imposed the moisture treatments. To inoculate plants, we grew the selected rhizobia strains in liquid TY for 48 h, diluted cultures to a standard optical density of 0.15 (OD_600_), and added 1 mL to each pot. We imposed moisture treatments every 2 days by watering plants assigned to the wet treatment to saturation while watering those assigned to the dry treatments to a 14% gravimetric water content. Unfortunately, there was a small outbreak of grasshopper herbivory in our greenhouse in the middle of this experiment. After eradicating the outbreak, we recorded which plants were grazed upon (approximately 5%) and quantified the damage, to be used as a factor in subsequent models.

Five weeks after the initiation of this experiment, we measured plant leaf number as a preliminary measure of plant growth. Eight weeks after the initiation of this experiment, we harvested both the above- and belowground biomass of plants over the course of 2 days.

Aboveground tissues were measured for their height, oven-dried at 75°C for 72 h, and then weighed. We gently washed belowground tissues of their soil and immediately stored them in a - 20°C freezer. As a proxy for rhizobia fitness, we dissected belowground tissues to count the number of root nodules (4–6), after which these belowground tissues were similarly dried and weighed.

#### Analyzing plant tissue nitrogen content

As rhizobia benefits are typically associated with nitrogen fixation, we evaluated the nitrogen content of a portion of plants inoculated with the evolved rhizobial strains. Tissues were ground and sent to the Stable Isotope Facility at the University of Wyoming (https://www.uwyo.edu/sif/), and analyzed for both total nitrogen and δ^15^N content. Given monetary and time costs associated with these analyses, and with ∼2000 plant samples, we only analyzed a small subset of samples. As one of our primary interests was in the impact of these rhizobia under drought, we first subset our samples to only those in the contemporary dry treatments. With minimal variation in plant health among those inoculated with rhizobia that evolved without a plant (see Results), these were removed as well. Amongst the remaining rhizobia strains, we randomly chose a single replicate from each strain, for a total of 160 samples.

#### Sequencing of rhizobial strains

To identify the bacterial genetic mechanisms underlying the evolved changes in rhizobial partner quality, we used long-read sequencing to build full genome assemblies for all strains evolved with a plant partner. With a high cost of sequencing, we focused specifically on this set of strains as minimal variation was found in those evolved without plant partners. Strains were sequenced by first growing each strain to a high density in liquid TY, after which we followed standard protocols using Zymo Quick-DNA HMW MagBead Kit (Irvine, CA, USA) to extract high molecular weight DNA from 1000 uL of sample. DNA was sent for PacBio HiFi long-read sequencing at the W. M. Keck center at the University of Illinois, where gDNAs were sheared with a Megaruptor 3 to an average fragment length of 10kb then converted to barcoded libraries with the SMRTBell Express Template Prep kit 3.0 and pooled in equimolar concentration. The pooled libraries were sequenced on 2 SMRTcell 8M on a PacBio Sequel II using the CCS sequencing mode and a 30hs movie time. Circular consensus sequence (CCS) analysis was done using SMRTLink V11.0 using the following parameters: ccs - -min-passes 3 --min-rq 0.99 lima --hifi-preset SYMMETRIC - -split-bam-named --peek-guess.

To generate de novo closed genomes for each evolved strain, we used the recommended workflow for Trycycler (7). Namely, we first filtered raw reads by length (above 1 kbp) and quality (top 95%) using Filtlong (8). Filtered reads were then subsampled in 12 independent subsets, from which we used three assembly methods, Flye (9), Hifiasm (10) and Raven (11) , to generate independent whole-genome assemblies for four read subsets each. Trycycler was then used to generate consensus genome according to these 12 assemblies. We determined the ancestry of each replicon in each strain using FastANI (12), calculating the average nucleotide identity (ANI) between all ancestor and all descendant genomes for each replicon. Ancestry for each descendant replicon was assigned to the ancestor replicon with the highest ANI.

We identified variants between the evolved strains and their ancestors by running sequences through two SNP (single nucleotide polymorphism) calling pipelines, each with their own strengths. We used the breseq pipeline, which was specifically designed for bacterial experimental evolution studies, as it can identify large insertions and deletions (13). We additionally used a custom Spine-Nucmer pipeline (https://github.com/Alan-Collins/Spine-Nucmer-SNPs) which, while unable to identify large insertions and deletions like breseq, we have found to be more effective in identifying point level mutations. From these strains, we identified some form of *de novo* variation introduced in 97 of the strains. This variation included 5 large deletions (the smallest ∼ 4800 bp in length), 1 instance of plasmid loss, and 665 individual SNPs. These SNPs, however, were not evenly distributed, 395 of these SNPs were distributed across just 2 strains.

#### Analysis of greenhouse data

We characterized shifts in rhizobia quality by building mixed-effects models for the 3 plant traits we measured representing aboveground plant growth: aboveground biomass, height, and leaf number. While ideally we would characterize partner quality by examining metrics of plant fitness, such as seed or fruit number, our target organism can take months to flower and produce seed, making proxies a necessary limitation here. We interpret these traits to be general measures of plant growth. These same traits have frequently been used in the legume-rhizobia literature (14–16). We built separate models for each contemporary watering environment. Our models included as fixed effects: the rhizobia’s historical moisture treatment (wet or dry), the rhizobia’s historic plant treatment (plant present or no plant), the historical nitrogen treatment (no nitrogen addition or nitrogen addition), and the level of herbivory. We included as random effects: greenhouse block and the individual strain.

We constructed models and evaluated the fixed effects using the *lme4* and *lmerTest* packages (17, 18) in the R statistical environment.

As parsing the effects of all 3 plant traits simultaneously can be difficult, we additionally merged these traits together into a single principal components (PC) axis. This axis represented 74% of the variation, and was positively correlated with all 3 traits. We interpret higher values here to represent higher quality rhizobia, and use this metric for downstream analyses. This was paired with a multivariate approach for these plant traits, using a constrained ordination with the rda function from the *vegan* R package (19). We included identical variables as the univariate models: contemporary watering environment, the rhizobia’s historical historic moisture treatment, the rhizobia’s historic plant treatment, and the rhizobia’s historical nitrogen treatment. To control for variation in the greenhouse, we inserted the Block and Herbivory terms into the Condition wrapper within the rda function, which partials out variation before fitting the terms of interest. While Strain must be included in the model to prevent pseudo-replication, this term could not be included in the Condition wrapper due to collinearity with the primary model terms. Following prior methods, we consequently used a modified permutation approach with the how function from the *permute* R package (20), restricting permutations so that replicates from the same Strain are kept together.

We additionally built similar linear mixed effects models for the two other traits measured, including belowground biomass and nodule number, as well as the root-shoot ratio, which represents the relative investment of the plant into belowground resource, which can be important in the context of plant nutrient and water scavenging. We chose not to incorporate these belowground traits into our multivariate proxies for partner quality as their connection to plant growth and fitness are less clear.

As our starting *Rhizobium* population was a mix of 28 strains with varying partner quality under wet and dry environments (Fig. S1), the observed differential phenotypes between treatments could be due to both shifts in strain frequencies as well as introduced *de novo* mutations. We address a portion of this variation by characterizing plant phenotypic variation within plants provided with evolved strains that could be mapped to the same ancestral background. We evaluated the comparison between strains adapted to low nitrogen, and either wet and dry environments by focusing on ancestral backgrounds that had multiple replicate evolved strains in these treatments. By focusing strains from the wet and dry, low nitrogen selection treatments, we could evaluate if the observed context dependent benefits that specifically emerged between the wet and dry, low nitrogen treatments could also be found within single strains. We built linear mixed effects models for each trait measured analogous to those described above, as well multivariate approaches incorporating all growth traits. Here though, the ancestral lineage was included as a random effect. We additionally used these strains to specifically evaluate tradeoffs into partner quality between environments. To appropriately compare the variation within each ancestral background, we mean centered the evolved strains’ partner quality across environments within their ancestral background. We then evaluated the tradeoff in the evolved benefits in partner quality with these strains by conducting a standard linear regression.

Similar to above, we compared variation in the evolved rhizobia’s partner quality, based on ancestral lineage, between the nitrogen treatments. As we observed the most significant drops in partner quality in the nitrogen evolved strains specifically when evolved in wet environments, and in contemporary wet environments, we focused our comparisons here. We conducted similar statistical analyses as reported above.

We evaluated potential contamination and the sterility of the treatments by comparing all plant traits as well as nodule number between live and sterile controls. Plants assigned to the sterile control, inoculated with no rhizobia, were on average significantly smaller in every plant health metric than those inoculated with live rhizobia across both the watering treatments (Figure SX). However, some of these sterile control plants did have a small degree of nodulation, despite not being provided with any inoculum and growing in sterile soils, indicating some degree of cross-contamination in the greenhouse. This contamination was minimal, with control plants producing 9% as many nodules as those inoculated with live rhizobia. As the experiment was randomized, this contamination was likely random with respect to treatment. These patterns of minimal, but present, contamination is common when working with rhizobia (1, 15, 21). We consequently use a similar interpretation as prior work of our results in the context of contamination – namely, while these low levels of contamination may minimize variation between treatments, as we still observed significant treatment variation (see Results), these treatments are likely strong drivers of rhizobia evolution. Our observed effect size estimates may, in fact, be underestimated in the face of this slight contamination. In all other analyses we excluded the sterile control plants.

#### Comparative genomics

The isolation frequency of ancestral lineages can be construed as a characterization of each strain’s fitness and population structure. We characterized how selective treatments impacted population structure by using chi-square statistics between each treatment combination. We additionally evaluated ancestral strain quality as a driver of population structure by correlating strain quality with isolation frequency, using a poisson distribution with the glmer function in R. We used our prior estimates of strain quality from both wet and dry environments as predictors here (see the section titled, **“Characterizing partner quality of the 28 ancestral strains”** below).

Beyond population structure, plasmid structure may be selected upon independent of the main chromosome. Our prior work has classified the pSyms of this group into 4 major clades, which are discordant from the main chromosome (22). Consequently, we characterized differentiation in population structure between selective treatments by specifically analyzing the phylogenetic history symbiotic plasmid. We used chi-square statistic to determine if pSym phylogenetic clades were differently abundant between each pairwise treatment combination.

To identify regions of genomic differentiation due to selective differences between treatments, we generated F_ST_ estimates between populations. We first created core genome alignments of each population using the Spine-Nucmer pipeline (see above). With the Fst.stats function in the *PopGenome* package (23), we then calculated a sliding window of Fst values across the core genome to investigate regions under selection. We then, using a custom script, tested if candidate regions with high F_ST_ (> 0.2) were genomic outliers, or alternatively, whether this genetic differentiation might simply be attributed to drift via Brownian motion. Briefly, ancestor strains were randomly sampled with replacement into two separate groups of 40 (simulating the descendant populations), then core alignments were created and F_ST_ values were calculated at the two candidate regions. This process was repeated 1000 times to generate distributions for each region of interest.

To examine the presence/absence difference of genes between selective treatments – which is missed in the above F_ST_ analyses based on core genome – we ran a custom presence/absence analysis. We first annotated all descendant strains with NCBI’s Prokaryotic Genome Annotation Tool (PGAP; (24)) and ran PIRATE (25) on the gff files of the separate populations to generate gene lists. The gene lists were then filtered for duplicates and then joined together in R with the *dplyr* package (26). We estimated if presence/absence variation represented significant differences by conducting Fisher’s exact tests for each unique gene category. Extracted *p-*values were adjusted for false discovery using an Benjamini-Hochberg approach (27).

#### Characterization of SNPS

To evaluate the potential contribution of *de novo* mutations to the observed phenotypes, we placed the genes in which SNPs occurred into putative functional bins based on annotation pipelines. Traditional experimental evolution studies might conduct a genome wide association study (GWAS) to identify specific elements underlying phenotypes.

This, however, is difficult here as while our data has a high level of replication in total number of strains isolated, these were mapped back to our ancestral backgrounds, with low within background replication. Moreover, across these ancestral backgrounds, there were large genomic alterations differentiating strains, with many SNPs occurring in regions not shared across all strains, making between-background comparisons difficult. While not a direct evaluation of the SNP contributions to phenotype, we instead evaluated the distribution of SNPs by putative function. SNPs occurring consistently in similar functions within a treatment may be the functions underlying phenotypic differences between treatments.

We first annotated the genomes of all ancestral strains. We passed all reference genomes through prodigal, a standard pipeline for predicting prokaryotic protein-coding genes (28). Genes output from this pipeline we then predicted the potential functions through use of InterProScan (29). Each SNP we observed could therefore be associated with a list of associated potential GO terms. We analyzed the distribution of SNPs based on their association with GO terms using multivariate methods. As addressed above, two of our strains were hypermutators. We removed these from this analysis, leaving 270 SNPs. We created a matrix representing the distribution of SNPs and their associated functions for each evolved strain: rows represented each strain and columns represented the observed GO term. This matrix was populated with 0s and 1s, with a 1 indicating the presence of a SNPs associated with that GO term for that strain. From this matrix we derived a distance matrix, based on Hamming distances between each strain. We used this distance matrix in a distance based RDA to evaluate similarity between treatments, with selective nitrogen and watering treatments predictor variables. From this RDA model, we extracted the GO terms with the strongest predictor of their distribution, indicating potential functional differences between treatments. Overall, our approach here is likely a conservative approach for evaluating the functional distribution of SNPs. Annotation pipelines are inherently imprecise, with a large fraction of the genes that our SNPs occurred had no associated functions, and therefore had to be excluded from analysis.

### Assessing nitrogen benefits of evolved rhizobial strains

We evaluated whether the observed context dependencies in partner quality across the low nitrogen, dry versus wet evolved strains was due to shifts in the efficiency of nitrogen fixation across environments. efficiency. To this end, we conducted a small, follow-up experiment with a subset of our evolved strains (5 high quality dry-adapted, no nitrogen strains and 5 high quality wet-adapted, no nitrogen strains), growing them in the greenhouse under variable watering conditions, and either with out without added nitrogen. If drought benefits, for example, were driven primarily through bacteria becoming more efficient at nitrogen fixation under drought, we might expect these benefits to disappear when an external source of nitrogen was added. The strains used here were chosen for being those with the highest estimates of partner quality in the wet and dry environments. Experimental methods followed those described above. Briefly, surface sterilized *T. repens* seeds were planted in sterile root-wash soil mix in sterilized cone-tainers. Plants were inoculated one week after seeding with one of our 10 rhizobia strains, or a sterile control. We amended pots designated to the nitrogen treatment with ammonium nitrate, with biweekly additions leading to a total of 0.03 g added over the course of the experiments.

This level was similar to that in agronomic settings and are presumed to have saturated plant nitrogen requirements. Plants were harvested at the 8-week mark, where we collected similar phenotypic data as described above, including height, above and belowground biomass, leaf number, and height. We assessed plant growth across all measured traits, as well as through multivariate approaches, as described above.

### Characterizing partner quality of the 28 ancestral lineages

We investigated variation in the partner quality of the 28 ancestral lineages of rhizobia across our two moisture environments (wet vs. dry) in a separate controlled greenhouse experiment. We followed similar methods as above, starting this experiment by filling sterilized cone-tainers with approximately 120 mL of root-wash mix. Soil was autoclaved (3 times in 1 h cycles, with 20 min. rests between cycles) shortly before the experiment to ensure the sterility of the soil. We then sprinkled 5 surface-sterilized seeds in each pot. For every treatment combination of rhizobia strain and moisture treatment, there were 15 replicates, for a total of 840 plants (28 rhizobia treatments x 2 moisture treatments x 15 replicates).

After seeding, we watered all pots every 2 days till saturation. Seven days after seeding, we inoculated plants with 1 mL of their designated rhizobia treatment, standardized to an optical density of 0.15 (OD_600_). One week after inoculation, we began the designated moisture treatments. We continued to water plants assigned to the wet treatment to saturation every 2 days, while we water plants assigned to the dry treatment to 14% gravimetric water content every 2 days. For the entirety of this experiment, plants were grown in the greenhouse on a 26°C/24°C day/night schedule, supplemented with 14 h of daily light. Ten weeks after the initiation of the experiment (and 7 since the start of the water treatments) we harvested both above- and belowground biomass of the plants over the course of 2 days. After harvest, we gently washed root tissues to remove soil particles. Aboveground tissue was measured for height and leaf number. All tissues were oven dried and then weighed. We counted the number of nodules on each root system as a proxy of rhizobia fitness. Similar to the above approaches, we compressed the aboveground traits to PCA to broadly evaluate growth along a single axis. We treat these multivariate compressions of plant growth as proxies for *Rhizobium* partner quality.

## SUPPLEMENTAL MATERIAL

**Fig. S1:**
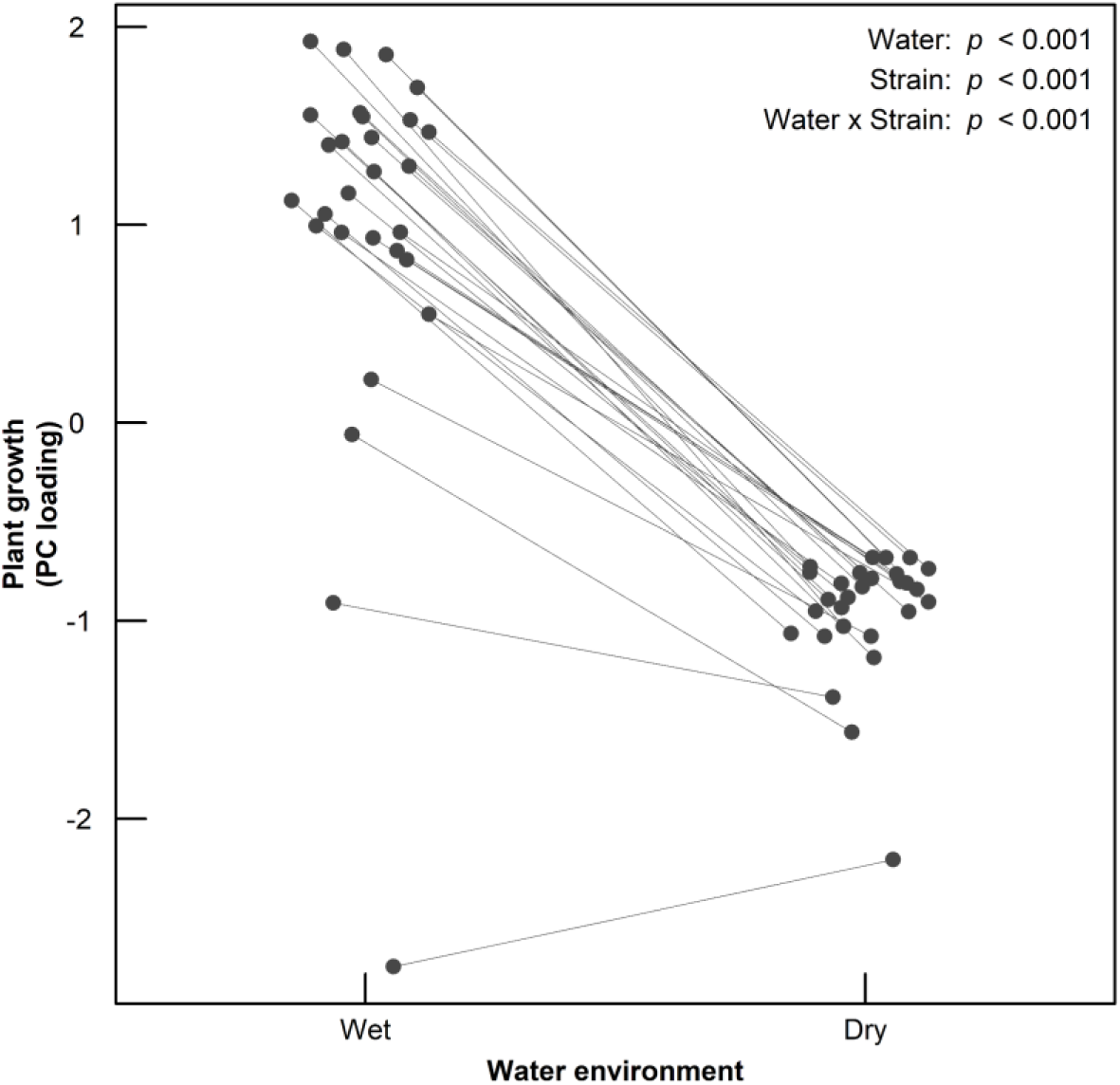
Our 28 ancestral *Rhizobium* lineages vary in partner quality across wet and dry environments. Partner quality here is characterized by a PC axis generated from multiple plant traits associated with growth, including aboveground biomass, leaf number, and height. We display *p* values representing a model describing the interaction in water and strain in the top right corner. Each point represents the marginal mean from this model for each distinct ancestral lineage. We emphasize that, as the data displayed here were generated from a separate greenhouse experiment as the experimentally evolved strains, these are not directly comparable with our evolved data. We exclude error bars from each strain to facilitate readability. Full details on data generation of the separate greenhouse experiment characterizing partner quality for these ancestral lineages can be found in the Supplemental Methods.

**Fig. S2:**
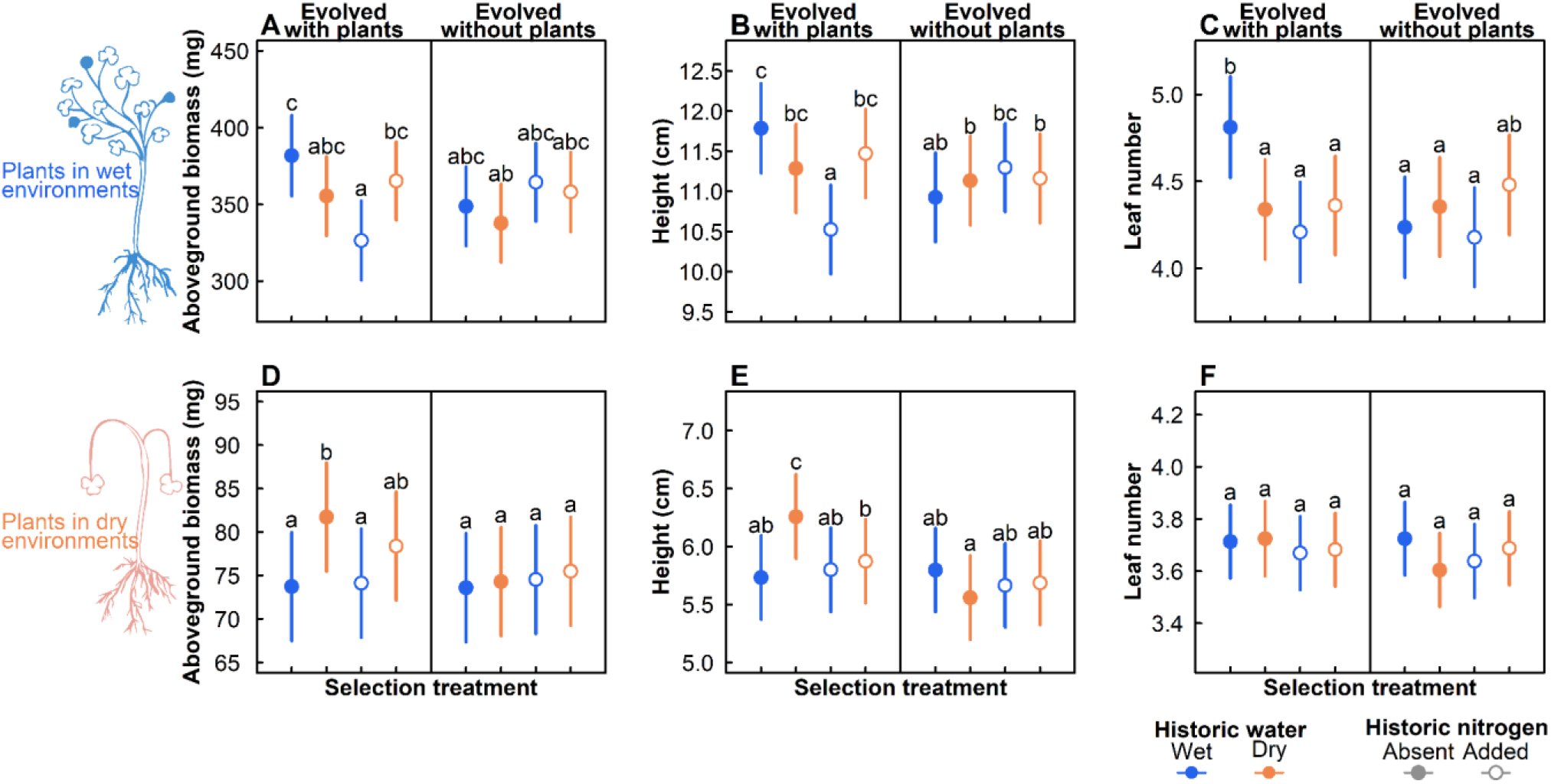
Evolutionary history of the experimentally evolved *Rhizobium* populations impacts plant aboveground traits in both contemporary wet (A-C) or contemporary dry (D-E) watering treatments. We display here aboveground biomass, height, and leaf number. The data displayed here represent those inputted into the PCA displayed in Fig. 1. We present estimated treatment marginal means, with bars representing 95% confidence intervals generated from the standard error. Lettering represents *post hoc* Tukey tests, with groups that differ in their lettering being significantly different from one another. Note the separate y-axes scales in top and bottom panels.

**Fig. S3:**
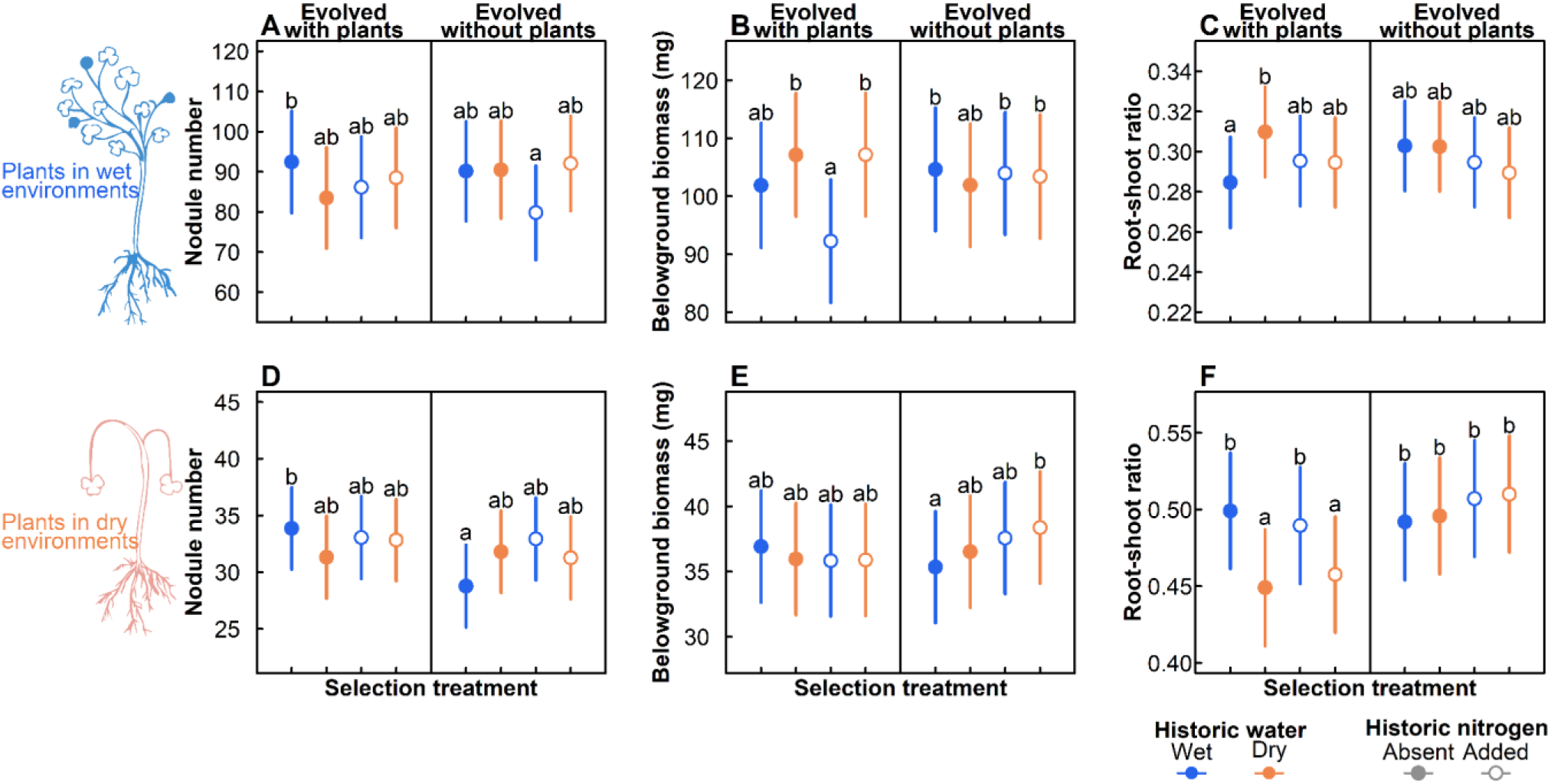
Evolutionary history of the experimentally evolved *Rhizobium* populations impacts plant aboveground traits in both contemporary wet (A-C) or contemporary dry (D-E) watering treatments. We display here nodule number, belowground biomass, and root-shoot ratio. We present estimated treatment marginal means, with bars representing 95% confidence intervals generated from the standard error. Lettering represents *post hoc* Tukey tests, with groups that differ in their lettering being significantly different from one another. Note the separate y-axes scales in top and bottom panels.

**Fig. S4:**
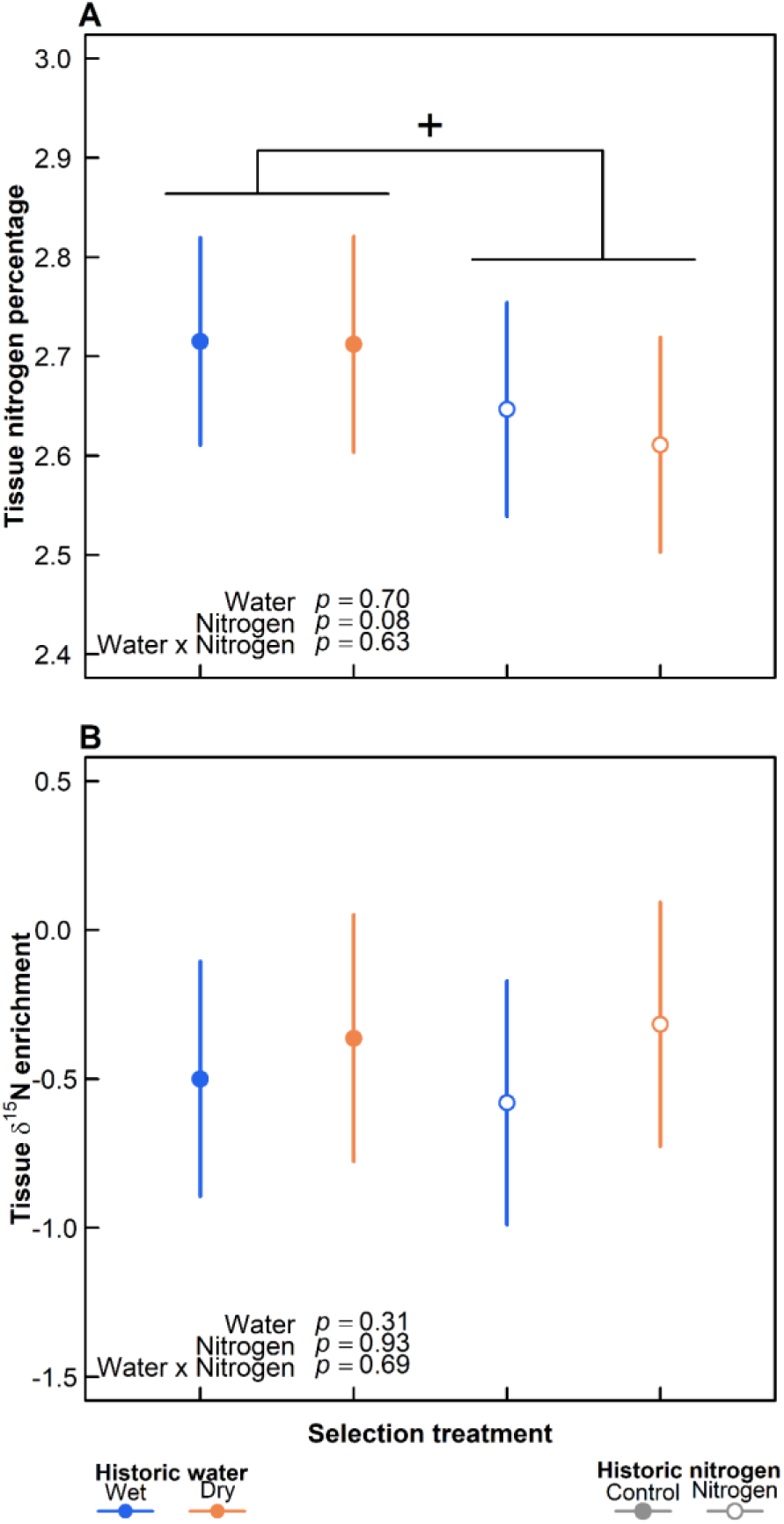
Evolutionary history of the experimentally evolved *Rhizobium* populations has a marginal impact on plant tissue nitrogen when grown in contemporary dry environments. We include both tissue nitrogen percentage **(A)** as well as tissue *δ*^15^N enrichment **(B)**, which has been used to quantify the proportion of nitrogen that derived from their rhizobial partners. These data only represent plants inoculated with strains that had evolved with a plant partner; those evolved without a plant partner were not analyzed due to cost constraints. Estimates represent marginal means, with bars representing 95% confidence intervals, extracted from a model including the historic water, historic nitrogen and their interactions. We display *p* values for the corresponding model terms in the bottom left corner for each panel.

**Fig. S5:**
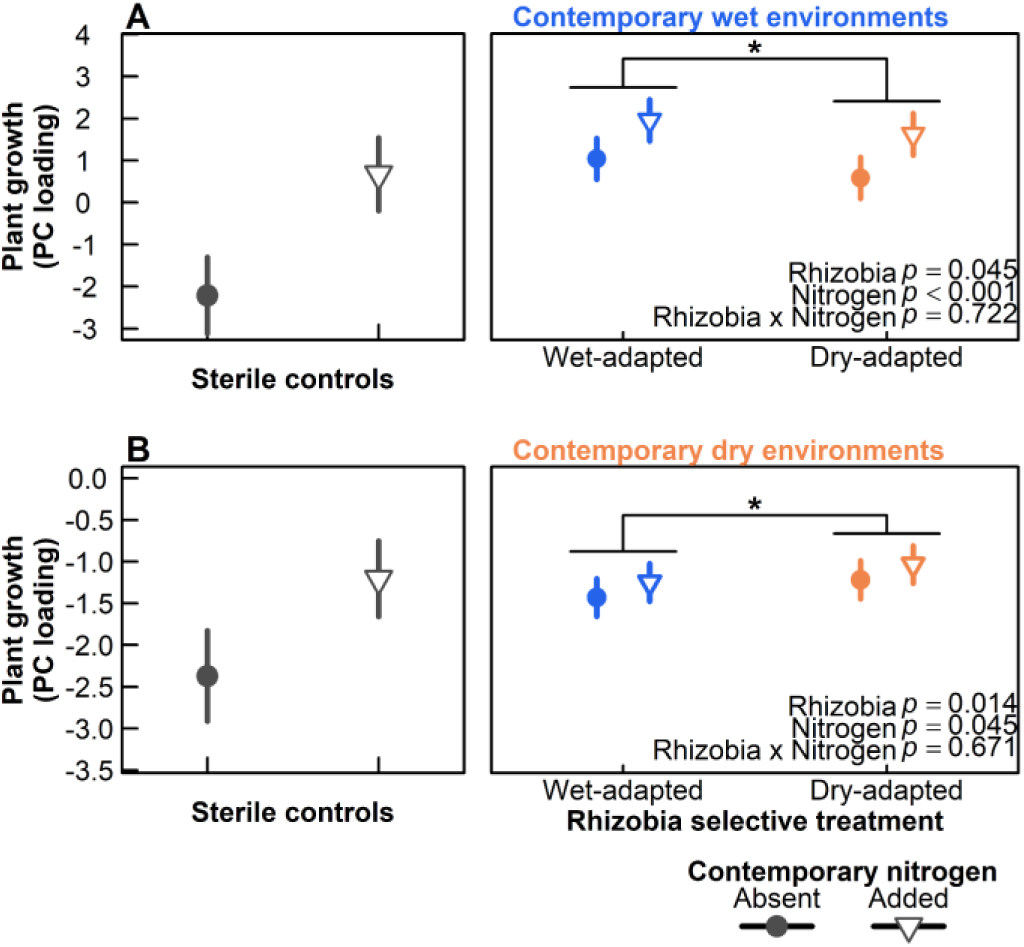
*Rhizobium* partner quality is maximized when evolved in a watering history that matched the contemporary watering condition, regardless of the contemporary nitrogen content under which we evaluated strain quality. We display here plant growth, extracted from a PC axis of aboveground plant traits including aboveground biomass, height, and leaf number, when grown with varying *Rhizobium* strain watering histories, while also varying the contemporary watering and nitrogen environments. All *Rhizobium* strains here evolved with a plant and under low nitrogen conditions, varying solely in their water history. We subset data to those plants grown in the contemporary wet **(A)** or contemporary dry **(B)** watering treatments. On the left side of each panel, we display the sterile controls to highlight the impacts of the nitrogen treatment. Estimates represent marginal means, with bars representing 95% confidence intervals, extracted from a model including the historic water, the contemporary nitrogen content, and their interactions. We include *p* values representing model terms in the bottom right corner. We note that these two figures have separate scales on their y-axis, as comparison within a group is difficult when placed on the same scale. Note these data were generated from a separate experiment as those displayed in the main text, and are therefore not directly comparable.

**Fig. S6:**
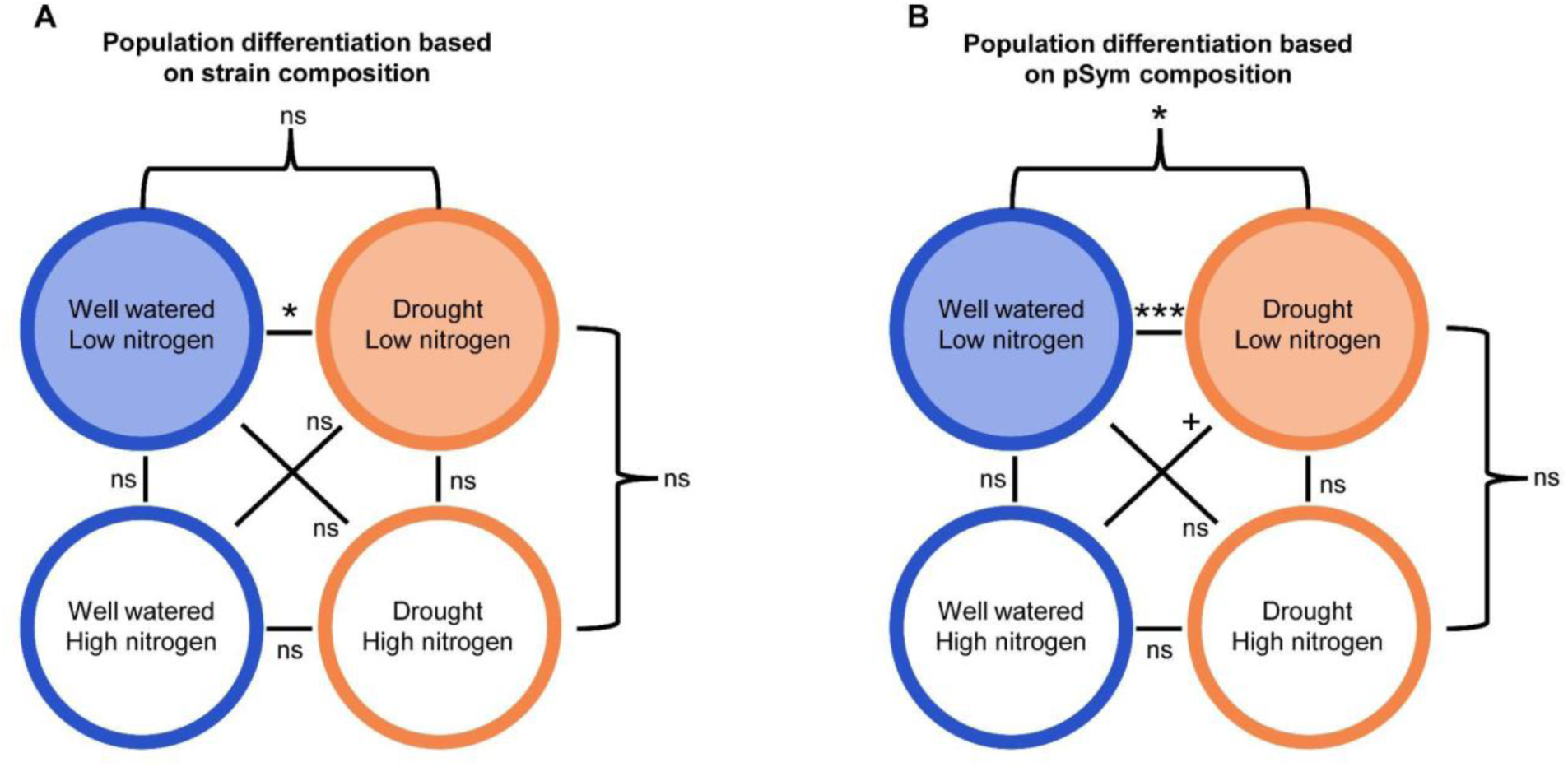
Historic selective treatments significantly impact the isolation frequency of the *Rhizobium* ancestral lineages. We display the pairwise comparisons between all treatment combinations wherein we compared isolation frequencies using χ^2^ tests. We compared isolation frequencies based both on identity within the 28 ancestral lineages (**A**), as well as across the 4 pSym types (**B**). We include *p* values from each comparison as labels with the following symbols: ns *p* > 0.1; + *p* ≤ 0.1; * *p* ≤ 0.05; ** *p* ≤ 0.01; *** *p* ≤ 0.001

**Fig. S7:**
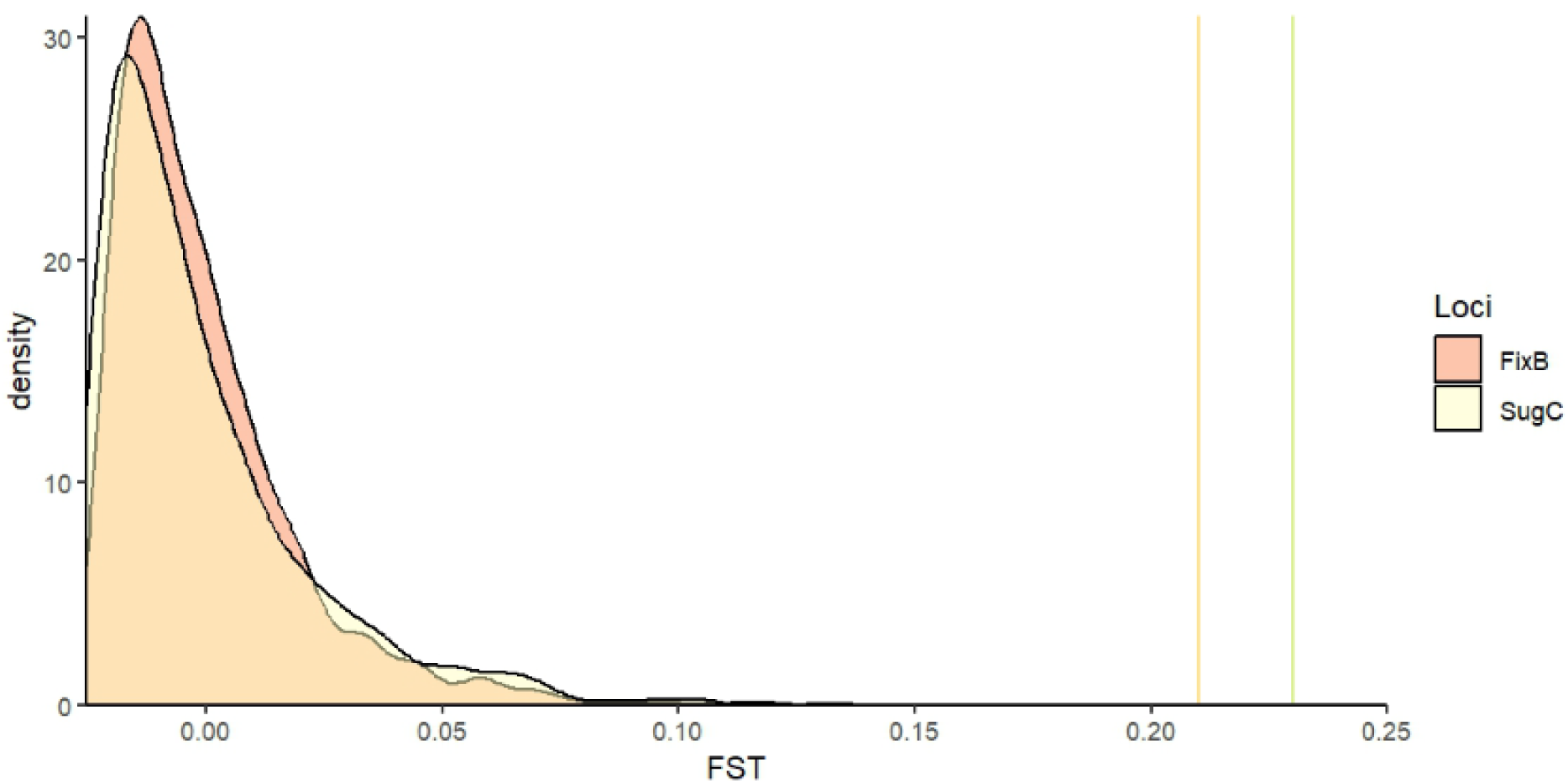
Observed F_ST_ along the fixB and sugC genes between watering treatments significantly differs in modeled evolution without selection. We display here the distribution of F_ST_ at these sites with random sampling of the ancestral strain population, representing evolution driven solely by drift. Observed F_ST_ for these sites are displayed as vertical lines at the far right of the figure.

**Fig S8:**
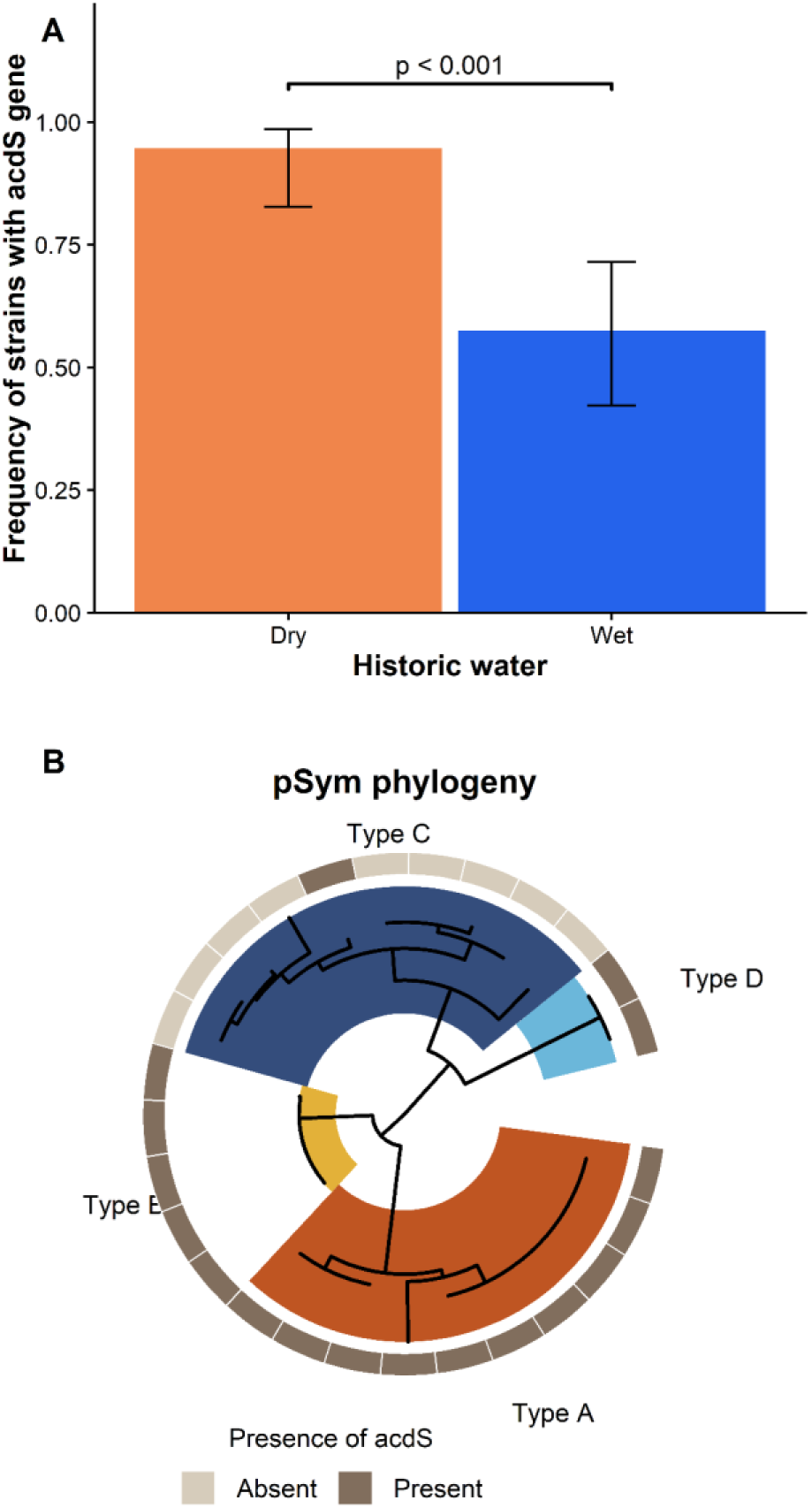
Significant variation in the presence of 1-aminocyclopropane-1-carboxylate deaminase genes (*acdS*) between watering treatments (A) and pSym types (B). We compare variation in the frequency of the *acdS* gene selective watering treatment, evolved without added nitrogen, using Fisher’s exact test. This variation is driven by pSym phylogeny, as the presence of this gene on the pSym is strongly correlated with phylogeny (seen in Fig. 3), being largely deplete type C pSyms.

**Fig. S9:**
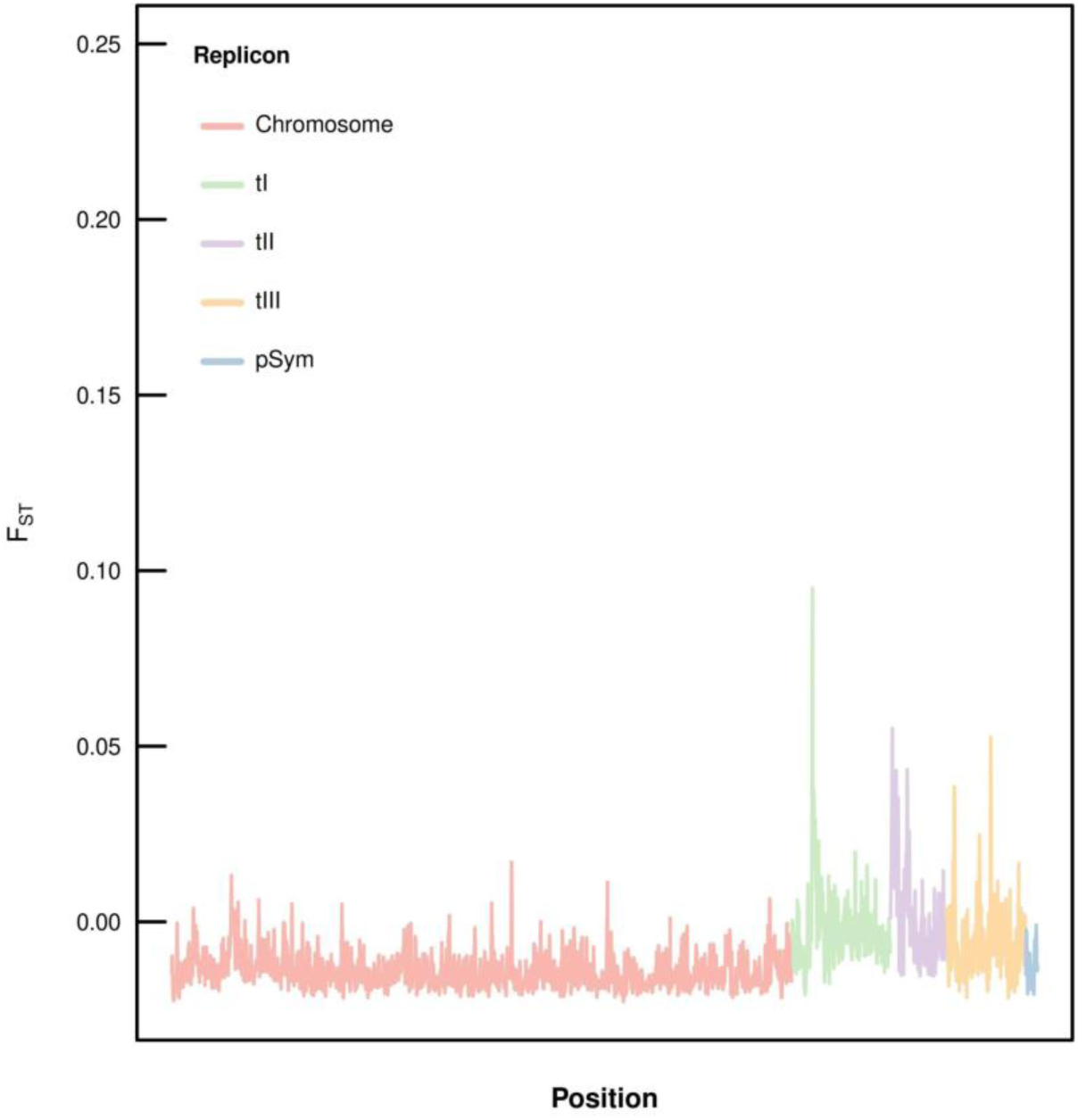
Minimal genomic variation between *Rhizobium* populations isolated between the two nitrogen treatments. We display F_ST_ estimates across the full genome between populations, with different replicons differentially colored, as coded in the legend in the top left corner. We specifically display here the comparison between no nitrogen added versus the nitrogen added treatments, in wet environments.

**Fig. S10:**
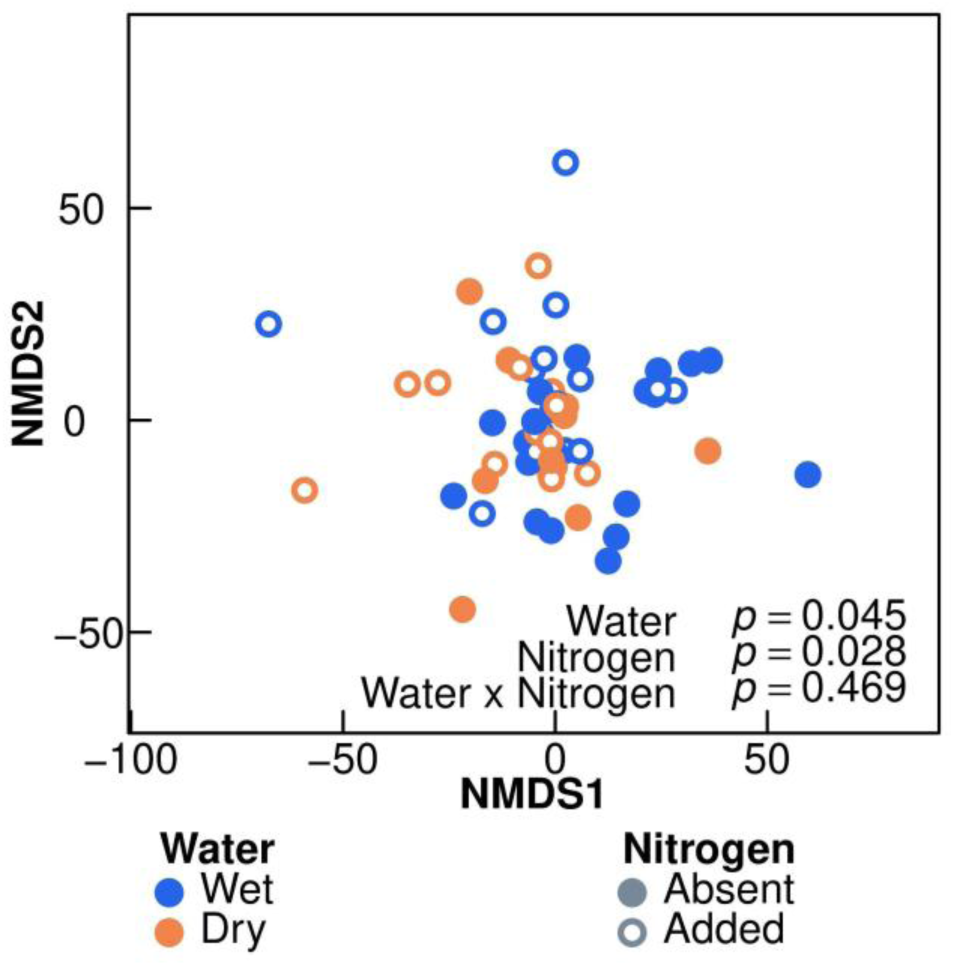
Strains’ evolutionary history is a significant predictor of the distribution of their SNPs, compared to their ancestral lineage, across gene annotations. We display here an NMDS for SNP distribution across genome annotations, with each point representing an individual evolved strain. Data are coded by the strains’ evolutionary history: the historical watering treatment (wet vs. dry, delineated by blue vs. orange coloring respectively) and the historical nitrogen treatment (unfertilized vs. fertilized, delineated by closed versus open circles respectively. See Table S9 for full model.

**Fig S11:**
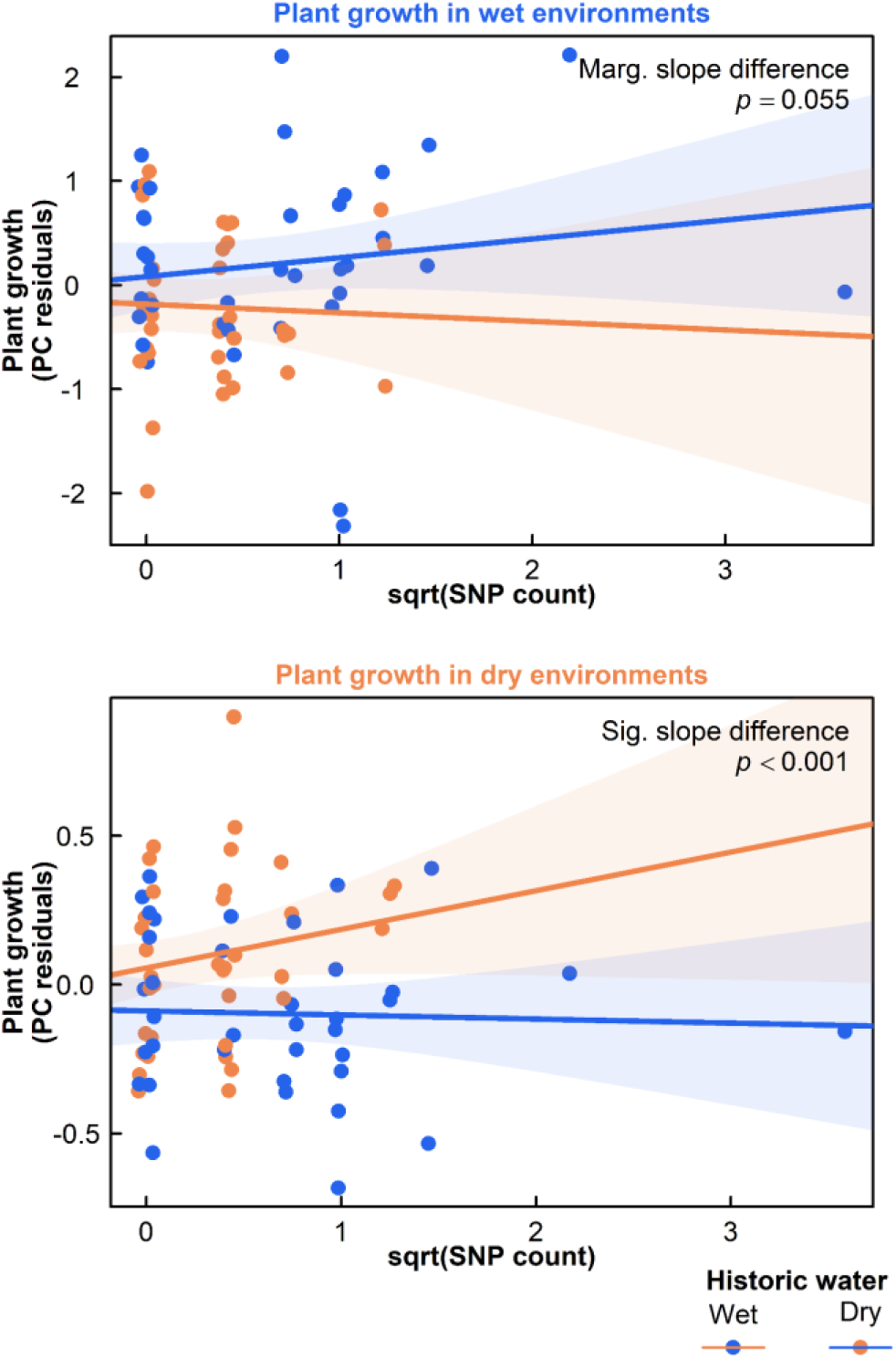
SNP count is correlated with increases in *Rhizobium* partner quality when in environments that matched the strains selective watering environment. Partner quality, extracted from the PC axis of aboveground plant growth traits (displayed in Fig. 1, S2, & S3), is standardized by extracting residual variation after ancestor lineage has been accounted for. Each point represents an individual evolved strain from those evolved without nitrogen addition, coded by their historical watering treatment: wet as blue, and dry as orange. We modeled partner quality as a function of the interacting historic watering treatments, and SNP count (square root transformed); the corresponding predictions for these models can also be seen here, including 95% prediction intervals. We include the *p* value for the interaction term in this model in the top right, representing the difference in slopes between watering treatments in each environment.

**Fig. S12:**
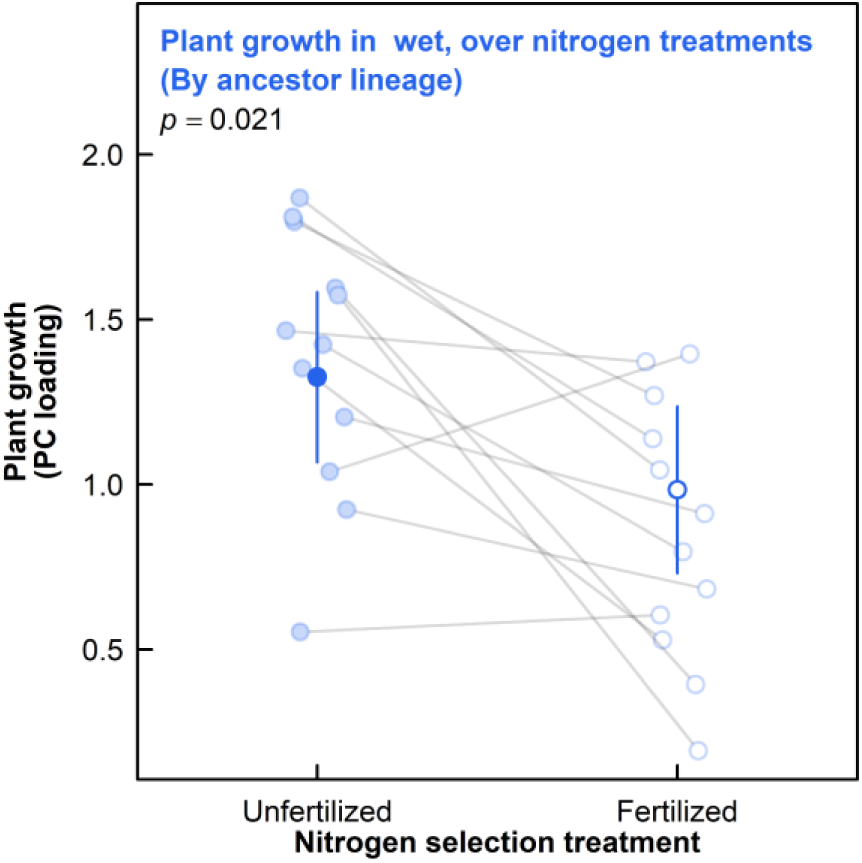
Nitrogen history of the experimentally evolved *Rhizobium* populations significantly impacts partner quality, even when controlling for ancestral lineage. These data are subset from those in Fig. 1, including only strains that map to ancestral backgrounds with isolation events in both unfertilized and fertilized selective treatments, from historic wet environments. We specifically show these strains plant growth in the contemporary wet environment. Semi-transparent points represent each ancestral lineage, while the solid point represents the group mean controlled for ancestral lineage. Growth benefits are characterized by the PCA of plant aboveground traits, and points are coded by strains’ historical nitrogen treatment, solid as no fertilizer and open as fertilizer added. A *p* value is included in the top left for this comparison.

**Fig S13:**
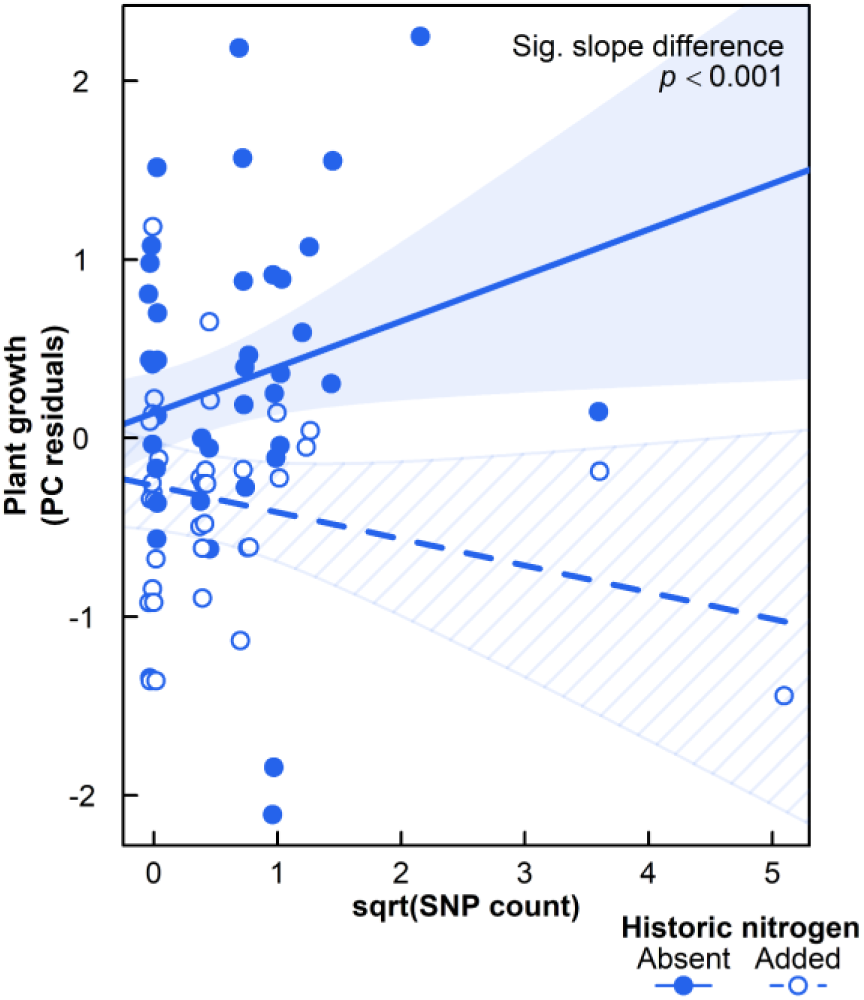
SNP count is correlated with decreases in *Rhizobium* partner quality when evolved in nitrogen fertilization treatments. Partner quality, extracted from the PC axis of aboveground plant growth traits (displayed in Fig. 1), is standardized by extracting residual variation after ancestor lineage has been accounted for. Each point represents an individual evolved strain from those evolved in wet environments, and coded by their historical nitrogen treatment: closed circles for unfertilized, and open circles for fertilized. We modeled partner quality as a function of the interacting historic nitrogen treatments, and SNP count (square root transformed); the corresponding predictions for these models can also be seen here, including 95% prediction intervals. We include the *p* value for the interaction term in this model in the top right, representing the difference in slopes between nitrogen treatments.

## SUPPLEMENTAL TABLES

**Table S1:** Evolutionary history of the experimentally evolved *Rhizobium* populations shapes their impact on a variety of plant traits in the contemporary wet environments. We display models in predicting various plant traits as a function of *Rhizobium* history, including the historic watering treatment (wet vs. dry), historic plant treatment (present vs. absent), historic nitrogen treatment (fertilized vs. unfertilized), and their interactions. Traits include above- and belowground biomass, leaf number, height, root-shoot ratio, and nodule number. These traits were modeled specifically using mixed effects models. We additionally include a multivariate model, built with an RDA, focusing specifically on traits directly associated with aboveground growth, including aboveground biomass, height, and leaf number. We display the associated *F* statistic for all model terms. We include the following symbology to indicate at level of significance: + *p* ≤ 0.1; * *p* ≤ 0.05; ** *p* ≤ 0.01; *** *p* ≤ 0.001.

| Model Terms | df | Biomass<br>(Aboveground) | Biomass<br>(Belowground) | Root-Shoot<br>Ratio | Height | Leaf number | Nodule number | Growth traits<br>(multivariate) |
| --- | --- | --- | --- | --- | --- | --- | --- | --- |
| HistoricWater | 1 | <0.01 | 2.20 | 0.61 | 0.73 | 0.07 | 0.21 | 0.19 |
| HistoricPlant | 1 | 0.22 | 0.23 | 0.04 | 0.86 | 1.53 | 0.05 | 0.82 |
| HistoricNitrogen | 1 | 0.06 | 0.61 | 1.38 | 1.27 | 1.81 | 0.57 | 1.01 |
| HistoricWater x HistoricPlant | 1 | 0.66 | 4.30* | 1.64 | 0.39 | 3.79<sup>+</sup> | 2.41 | 1.38 |
| HistoricWater x HistoricNitrogen | 1 | 3.65<sup>+</sup> | 1.10 | 1.75 | 3.42<sup>+</sup> | 4.55* | 3.29<sup>+</sup> | 4.45* |
| HistoricPlant x HistoricNitrogen | 1 | 5.46* | 0.85 | 0.49 | 6.08* | 2.90<sup>+</sup> | 0.33 | 4.89** |
| HistoricWater x HistoricPlant x<br>HistoricNitrogen | 1 | 2.63 | 0.44 | 0.79 | 9.02** | 1.36 | <0.01 | 5.18** |

**Table S2:**
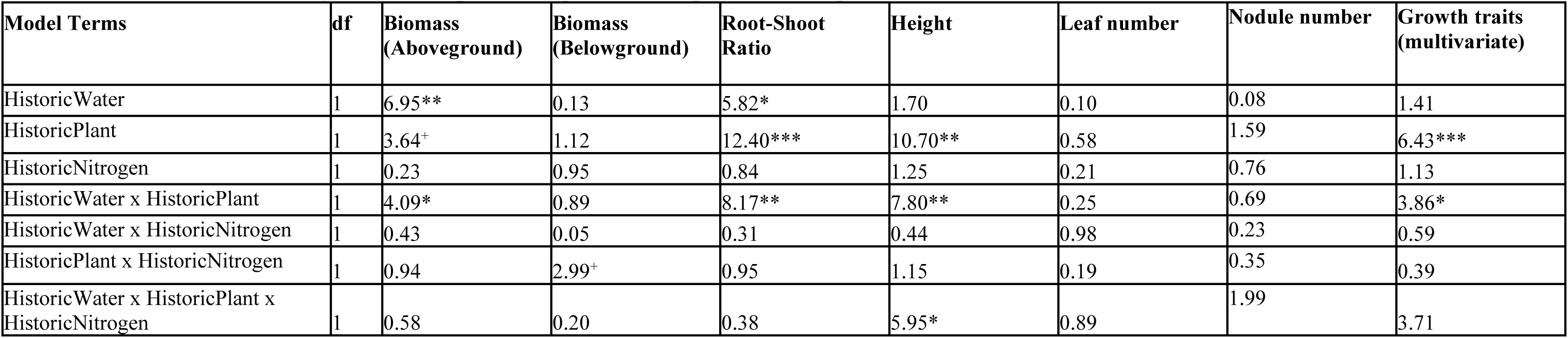
Evolutionary history of the experimentally evolved *Rhizobium* populations shapes their impact on a variety of plant traits in the contemporary dry environments. We display models in predicting various plant traits as a function of *Rhizobium* history, including the historic watering treatment (wet vs. dry), historic plant treatment (present vs. absent), historic nitrogen treatment (fertilized vs. unfertilized), and their interactions. Traits include above- and belowground biomass, leaf number, height, root-shoot ratio, and nodule number. These traits were modeled specifically using mixed effects models. We additionally include a multivariate model, built with an RDA, focusing specifically on traits directly associated with aboveground growth, including aboveground biomass, height, and leaf number. We display the associated *F* statistic for all model terms. We include the following symbology to indicate at level of significance: + *p* ≤ 0.1; * *p* ≤ 0.05; ** *p* ≤ 0.01; *** *p* ≤ 0.001.

| Model Terms | df | Biomass<br>(Aboveground) | Biomass<br>(Belowground) | Root-Shoot<br>Ratio | Height | Leaf number | Nodule number | Growth traits<br>(multivariate) |
| --- | --- | --- | --- | --- | --- | --- | --- | --- |
| HistoricWater | 1 | 6.95** | 0.13 | 5.82* | 1.70 | 0.10 | 0.08 | 1.41 |
| HistoricPlant | 1 | 3.64<sup>+</sup> | 1.12 | 12.40*** | 10.70** | 0.58 | 1.59 | 6.43*** |
| HistoricNitrogen | 1 | 0.23 | 0.95 | 0.84 | 1.25 | 0.21 | 0.76 | 1.13 |
| HistoricWater x HistoricPlant | 1 | 4.09* | 0.89 | 8.17** | 7.80** | 0.25 | 0.69 | 3.86* |
| HistoricWater x HistoricNitrogen | 1 | 0.43 | 0.05 | 0.31 | 0.44 | 0.98 | 0.23 | 0.59 |
| HistoricPlant x HistoricNitrogen | 1 | 0.94 | 2.99<sup>+</sup> | 0.95 | 1.15 | 0.19 | 0.35 | 0.39 |
| HistoricWater x HistoricPlant x<br>HistoricNitrogen | 1 | 0.58 | 0.20 | 0.38 | 5.95* | 0.89 | 1.99 | 3.71 |

**Table S3:**
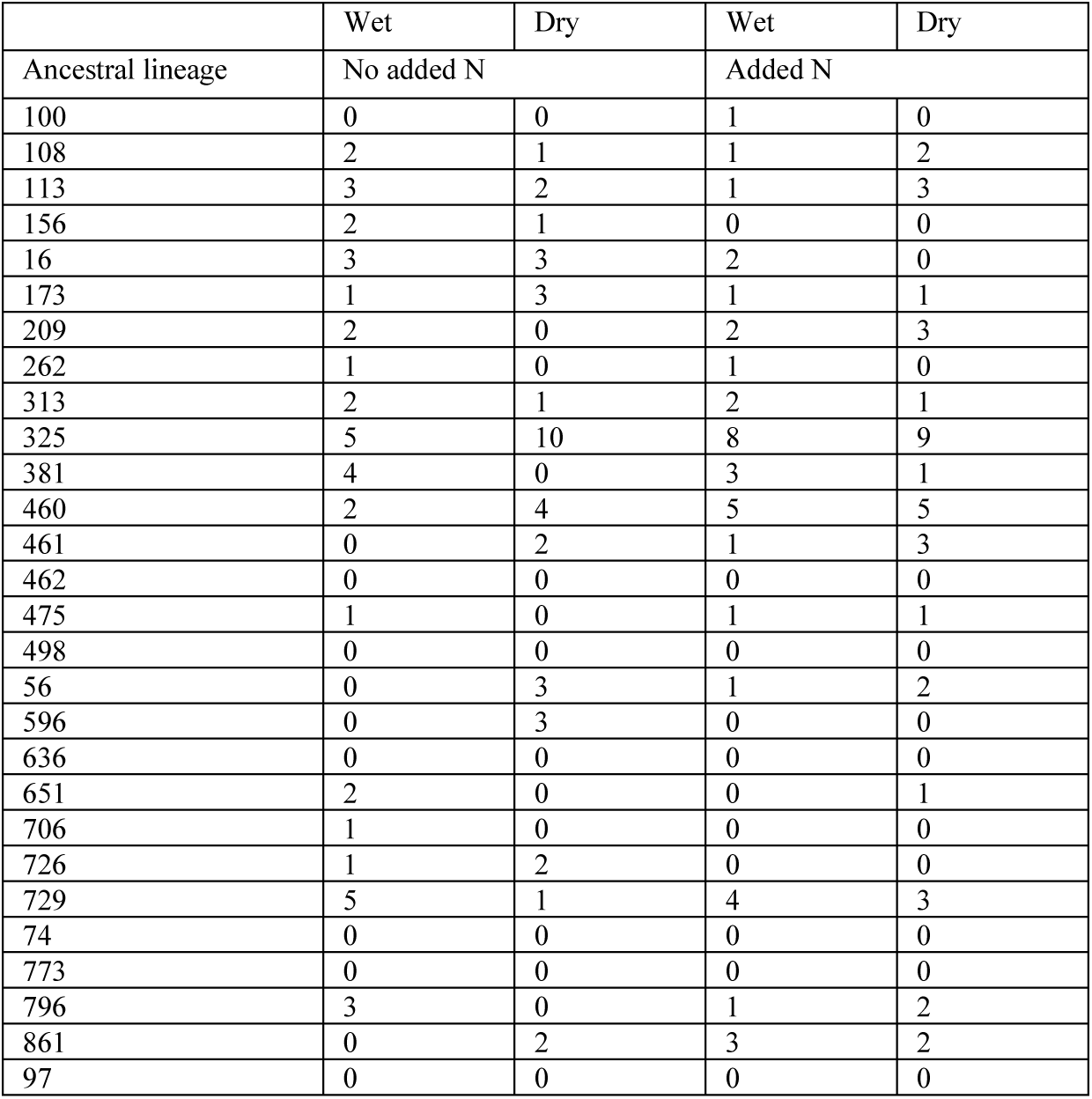
Isolation frequency of evolved strains mapped back to the original 28 ancestral lineages, broken down by selective treatment. Note, these are solely the treatments evolved with a plant present, as the plant absent treatments were not sequenced.

**Table S4:** Partner quality from matched environments predicts isolation frequency in a given watering environment with partner quality from the same environment. Comparisons between two models predicting the reisolation frequency of *Rhizobium* ancestral lineages based on the ancestors’ partner quality across the two watering environments. Our proxy for partner quality here is the PC axis combining aboveground biomass, height, and leaf count, identical to that seen in Fig. S1. The reported models are both generalized linear mixed-effects models with poisson distributions, incorporating the ancestral lineage as a random effect. Ancestral isolation frequency in wet environments is much better predicted by their partner quality in wet, and isolation frequency in dry environments by their partner quality in dry (as opposed to mismatched environments).We present the R^2^ for these models, as well as Akaike weights, derived from AIC, which represents the probability a given model is the best model within the set We incorporate the following symbology in the R^2^ column to indicate at level of significance for the model: + *p* ≤ 0.1; * *p* ≤ 0.05; ** *p* ≤ 0.01; *** *p* ≤ 0.001.

| Model | $R^2$ | Probability of best model (Akaike weights) |
| --- | --- | --- |
| Isolation frequency~Partner quality from matched environment | 0.37** | 0.97 |
| Isolation frequency~Partner quality from mismatched environment | 0.13<sup>+</sup> | 0.03 |

**Table S5:**
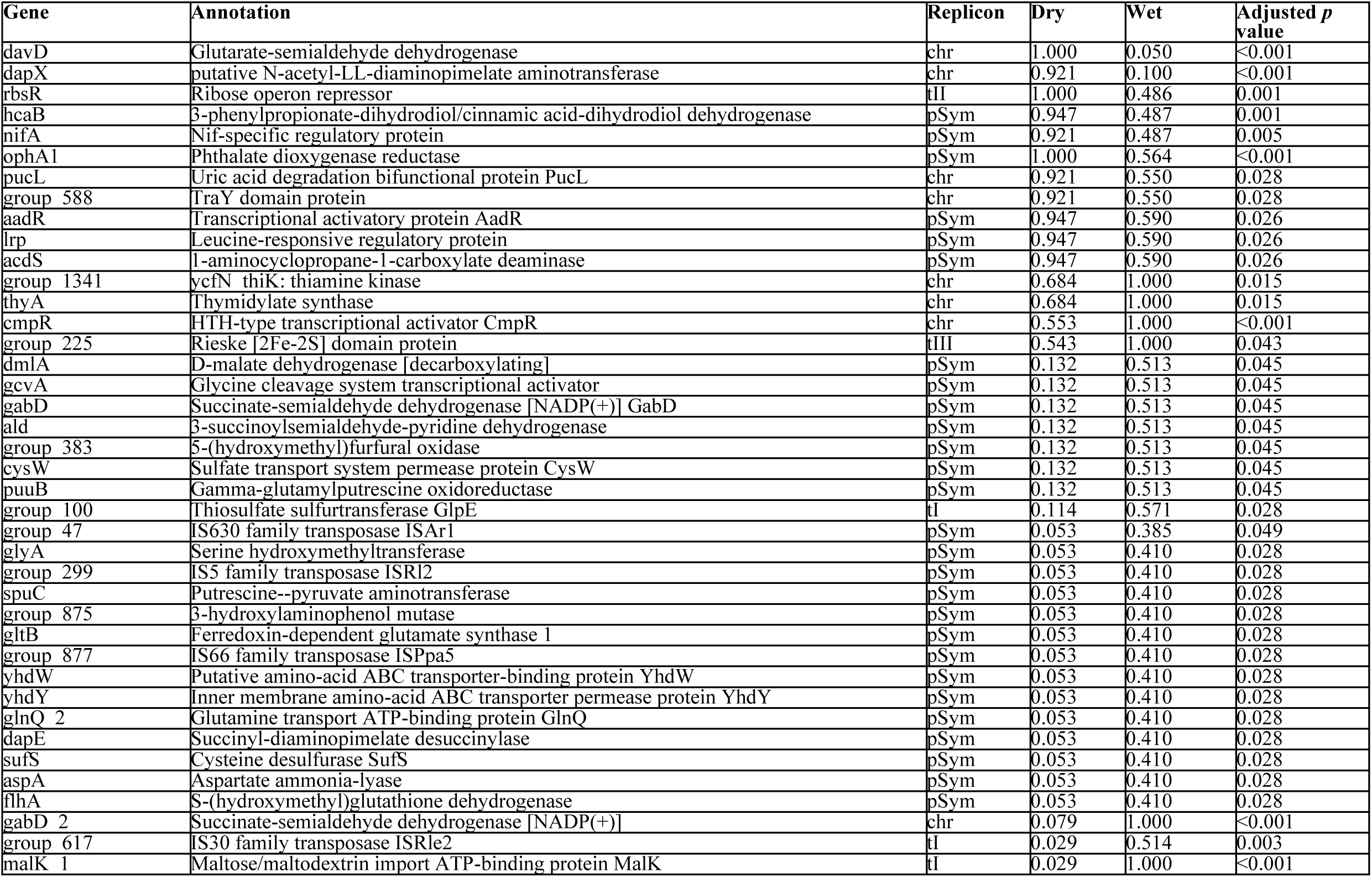
Selective watering history of experimentally evolved *Rhizobium* populations generates significant differences in gene abundances between watering treatments. We highlight the 40 genes with significantly different presence/absence variation (PAV) between the wet and dry, low nitrogen adapted populations. We include gene product name, functional annotation, the replicon the gene is on, the frequency in each treatment, as well as a *p* value (adjusted for multiple testing) from the Fisher’s test comparing treatments. We use shorthand in the replicon column, following terminology for *Rhizobium* plasmids described in Vereau-Gorbitz et al. 2025: chr-main chromosome, tI-Type I plasmid, tII-Type II plasmid, tIII-Type III plasmid, & pSym-the symbiotic plasmid.

**Table S6:**
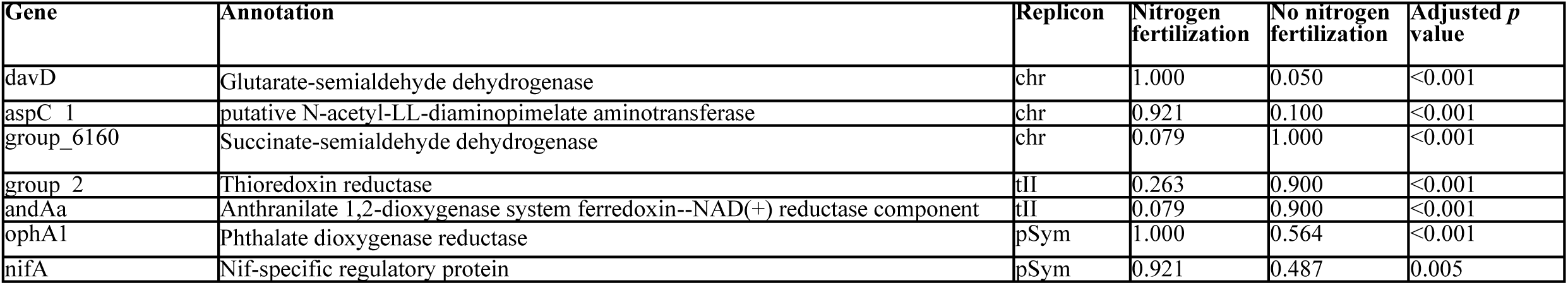
Selective nitrogen history of experimentally evolved *Rhizobium* populations generates significant differences in gene abundances between nitrogen treatments. We highlight the 5 genes with significantly different presence/absence variation (PAV) between the nitrogen addition and absent, wet adapted populations. We include gene product name, functional annotation, the replicon the gene is on, the frequency in each treatment, as well as a *p* value (adjusted for multiple testing) from the Fisher’s test comparing treatments. We use shorthand in the replicon column, following terminology for *Rhizobium* plasmids described in Vereau-Gorbitz et al. 2025: chr-main chromosome, tI-Type I plasmid, tII-Type II plasmid, tIII-Type III plasmid, & pSym-the symbiotic plasmid.

**Table S7:**
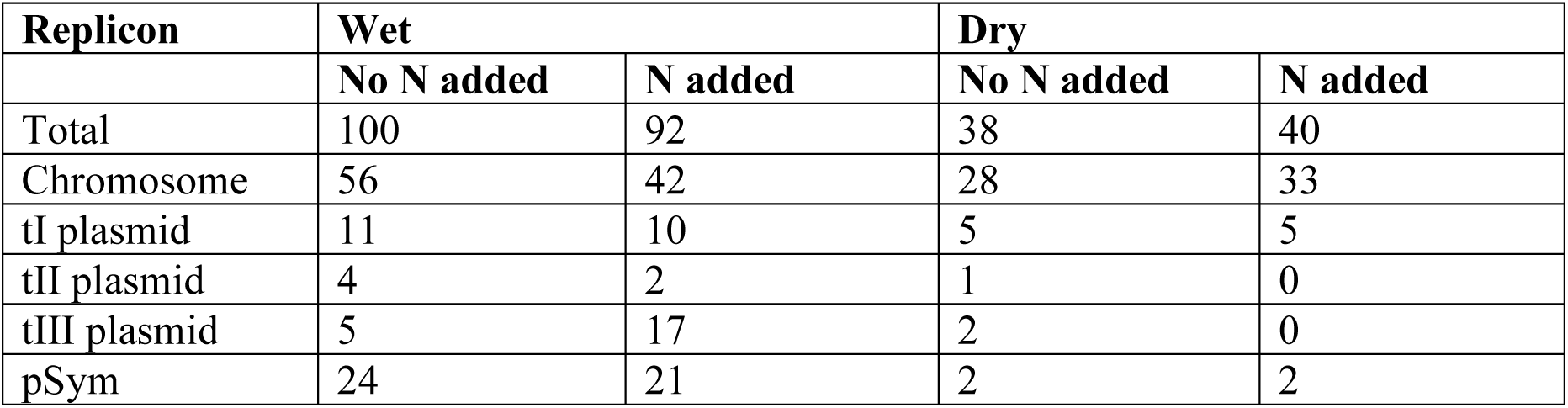
Total number of SNPs observed in our evolved strains, as compared to their ancestor’s genome. While the first line displays SNP counts for the full bacterial genome, we additionally break these counts down by each replicon.

**Table S8:** Rate of SNP occurrence observed in our evolved strains, as adjusted for sample and replicon size. Data presented here are directly related to the frequencies observed in Table S7, adjusting for the size and frequency of the replicon.

| Replicon | Wet |  | Dry |  |
| --- | --- | --- | --- | --- |
|  | No N added | N added | No N added | N added |
| Total | $3.43 \times 10^{-7}$ | $3.20 \times 10^{-7}$ | $1.30 \times 10^{-7}$ | $1.58 \times 10^{-7}$ |
| Chromosome | $2.86 \times 10^{-7}$ | $2.21 \times 10^{-7}$ | $1.44 \times 10^{-7}$ | $1.69 \times 10^{-7}$ |
| tI plasmid | $2.76 \times 10^{-7}$ | $2.39 \times 10^{-7}$ | $1.21 \times 10^{-7}$ | $1.19 \times 10^{-7}$ |
| tII plasmid | $1.99 \times 10^{-7}$ | $9.75 \times 10^{-8}$ | $4.88 \times 10^{-8}$ | 0 |
| tIII plasmid | $2.42 \times 10^{-7}$ | $8.12 \times 10^{-7}$ | $9.62 \times 10^{-8}$ | 0 |
| pSym | $1.57 \times 10^{-6}$ | $1.41 \times 10^{-6}$ | $1.31 \times 10^{-7}$ | $1.31 \times 10^{-7}$ |

**Table S9:**
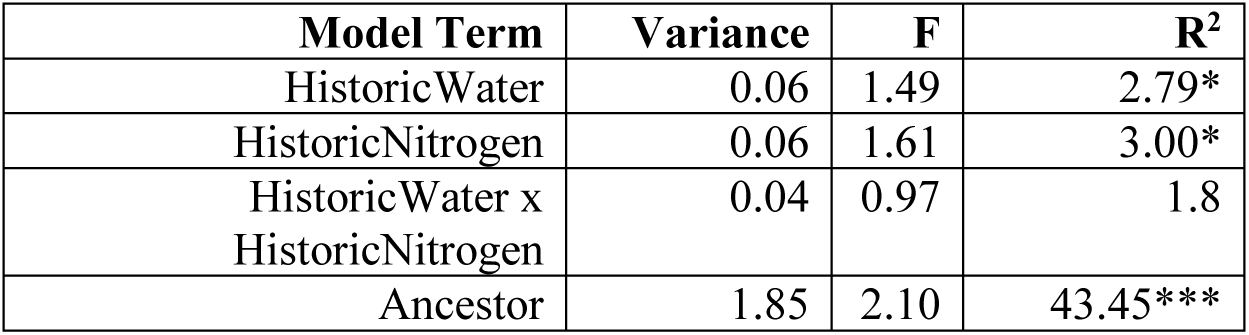
Evolutionary history is a significant predictor of the distribution of SNPs based on their locations within gene annotation categories. This table specifically represents an RDA model, with GO terms and their abundance of SNPs within each term as response variables. This Table corresponds to data displayed in Fig. S10, and we include the following symbology to indicate at level of significance: + p ≤ 0.1; * p ≤ 0.05; ** p ≤ 0.01; *** p ≤ 0.001.

**Table S10:** Historic watering history of the experimentally evolved *Rhizobium* populations impacts their partner quality in contemporary wet environments, even within ancestral lineages. These are model outputs for all plant traits measured and subsets of those reported in Table S1, while critically accounting for the ancestral lineages. Specifically, these models include strains evolved with a plant, with no nitrogen addition. We include ancestral lineage as a random effect to control for this variation. Identical traits to those reported in Table S1 are used. This Table corresponds to Fig. 4A. We display the associated F statistic for all model terms. Terms with a p value less than 0.1 are bolded for emphasis, and we include the following symbology to indicate at level of significance: + p ≤ 0.1; * p ≤ 0.05; ** p ≤ 0.01; *** p ≤ 0.001.

| Model terms | df | Biomass<br>(Aboveground) | Biomass<br>(Belowground) | Root-Shoot<br>Ratio | Height | Leaf number | Nodule number | Growth traits<br>(multivariate) |
| --- | --- | --- | --- | --- | --- | --- | --- | --- |
| HistWater | 1 | 1.05 | <0.01 | 2.77+ | 2.11 | 6.03* | 0.22 | 4.21* |
| Ancestor | 3 | 2.03 | 1.05 | 1.40 | 0.68 | 1.86 | 0.66 | 1.20 |
| HistWater x Ancestor | 3 | 0.45 | 0.78 | 0.65 | 0.54 | 0.28 | 2.06 | 0.21 |

**Table S11:** Historic watering history of the experimentally evolved *Rhizobium* populations impacts their partner quality in contemporary dry environments, even within ancestral lineages. These are model outputs for all plant traits measured and subsets of those reported in Table S2, while critically accounting for the ancestral lineages. Specifically, these models include strains evolved with a plant, with no nitrogen addition. We include ancestral lineage as a random effect to control for this variation. Identical traits to those reported in Table S2 are used. This Table corresponds to Fig. 4B. We display the associated F statistic for all model terms. Terms with a p value less than 0.1 are bolded for emphasis, and we include the following symbology to indicate at level of significance: + p ≤ 0.1; * p ≤ 0.05; ** p ≤ 0.01; *** p ≤ 0.001.

| Model terms | df | Biomass<br>(Aboveground) | Biomass<br>(Belowground) | Root-Shoot<br>Ratio | Height | Leaf number | Nodule number | Growth traits<br>(multivariate) |
| --- | --- | --- | --- | --- | --- | --- | --- | --- |
| HistWater | 1 | 5.96* | 0.03 | 3.36+ | 7.17** | 1.99 | 0.033 | 6.96* |
| Ancestor | 3 | 0.82 | 3.03* | 0.90 | 0.23 | 1.50 | 0.20 | 1.26 |
| HistWater x Ancestor | 3 | 0.51 | 0.76 | 0.93 | 0.30 | 1.24 | 0.52 | 0.67 |

**Table S12:**
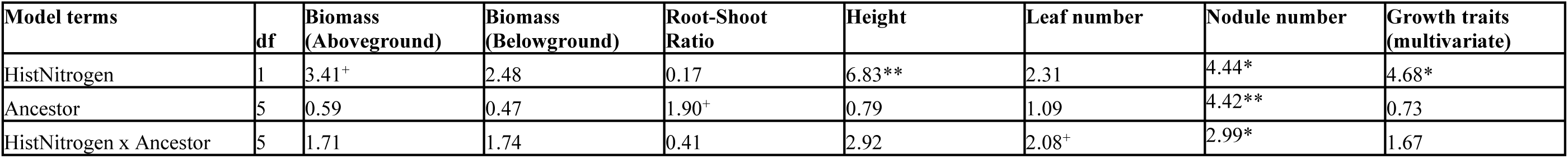
Historic nitrogen history of the experimentally evolved *Rhizobium* populations impacts their partner quality, even within ancestral lineages. These are model outputs for all plant traits measured and subsets of those reported in Table S1, while critically accounting for the ancestral lineages. Specifically, these models include strains evolved with a plant, under wet conditions. We include ancestral lineage as a random effect to control for this variation. Identical traits to those reported in Table S1 are used. This Table corresponds to Fig. S12. We display the associated F statistic for all model terms. Terms with a p value less than 0.1 are bolded for emphasis, and we include the following symbology to indicate at level of significance: + p ≤ 0.1; * p ≤ 0.05; ** p ≤ 0.01; *** p ≤ 0.001.

| Model terms | df | Biomass<br>(Aboveground) | Biomass<br>(Belowground) | Root-Shoot<br>Ratio | Height | Leaf number | Nodule number | Growth traits<br>(multivariate) |
| --- | --- | --- | --- | --- | --- | --- | --- | --- |
| HistNitrogen | 1 | 3.41<sup>+</sup> | 2.48 | 0.17 | 6.83** | 2.31 | 4.44* | 4.68* |
| Ancestor | 5 | 0.59 | 0.47 | 1.90<sup>+</sup> | 0.79 | 1.09 | 4.42** | 0.73 |
| HistNitrogen x Ancestor | 5 | 1.71 | 1.74 | 0.41 | 2.92 | 2.08<sup>+</sup> | 2.99* | 1.67 |

